# Identification of bridgin, an unconventional linker, connects the outer kinetochore to centromeric chromatin

**DOI:** 10.1101/816199

**Authors:** Shreyas Sridhar, Tetsuya Hori, Reiko Nakagawa, Tatsuo Fukagawa, Kaustuv Sanyal

## Abstract

The microtubule-binding outer kinetochore is linked to centromeric chromatin through the inner kinetochore CENP-C^Mif2^, CENP-T^Cnn1^, and CENP-U^Ame1^ pathways. These are the only known kinetochore linker proteins across eukaryotes. Linker proteins are structurally less conserved than their outer kinetochore counterparts. Here, we demonstrate the recurrent loss of most inner kinetochore CCAN, including certain linker proteins during evolution in the fungal phylum of Basidiomycota. By studying the kinetochore interactome, a previously undescribed linker protein, bridgin was identified in the basidiomycete *Cryptococcus neoformans*, a human fungal pathogen*. In vivo and in vitro* functional analyses of bridgin reveal that it binds to the outer kinetochore and centromere chromatin simultaneously to ensure accurate kinetochore-microtubule attachments. Unlike known linker proteins, bridgin is recruited by the outer kinetochore. Homologs of bridgin were identified outside fungi. These results showcase a divergent strategy, with a more ancient origin than fungi, to link the outer kinetochore to centromeric chromatin.

---

Accurate chromosome segregation ensures faithful transmission of the genetic material to the progeny. The kinetochore is a multi-complex protein network that assembles on the centromere of each chromosome^1–4^ and is attached to the spindle microtubules for accurate chromosome segregation ^5^. Components involved in error correction mechanisms and the spindle assembly checkpoint (SAC) are recruited at kinetochores to ensure bi-orientation of sister chromatids in mitosis^6–9^. The inner kinetochore is composed of CENP-A^10–12^ (centromeric Histone H3 variant) and the 16-member (in vertebrates) constitutive centromere associated network (CCAN)^13–16^. The outer kinetochore members of the KMN (KNL1, Mis12, and Ndc80 complexes) network^1, 17^ are recruited to CCAN to form the kinetochore ensemble in various model systems including budding yeast and vertebrate cells ^18^. Additional components such as the three-member Ska1 complex in vertebrates and the 10-member Dam1 complex (Dam1C) in fungi localize to the outer kinetochore to ensure accurate kinetochore-microtubule interactions^19–24^.

The Ndc80 complex (Ndc80C) of the KMN network directly binds to spindle microtubules ^25–27^. CCAN proteins, CENP-C^Mif2^, and CENP-T^Cnn1^ have been shown to bridge centromeric chromatin with the KMN network independently^28–32^. Additionally, CENP-U^Ame1^ functions as a linker in budding yeast^33, 34^. CENP-C^Mif2^ and CENP-T^Cnn1^, through their amino-termini (N), interact with the Mis12 complex (Mis12C) and the Ndc80C respectively, while their carboxy(C)-termini interact with centromeric chromatin^35–37^. An extended unstructured region separates the N and C termini in both CENP-C^Mif2^ and CENP-T^Cnn1^. CENP-U^Ame1^ also binds to Mis12C and CENP-A^Cse4^ simultaneously to ensure a linker function in budding yeast^33, 34^. Although these linker pathways are essential for accurate chromosome segregation, their architectural dependence varies across species^29, 30, 32^.

With the availability of genome sequencing data of a large number of organisms and improved bioinformatic tools, it has become evident that the KMN network components are highly conserved across eukaryotes^38, 39^. Further, organisms with varying kinetochore compositions have been described, and such variations predominantly emerge due to alterations in the inner kinetochore composition^22, 40–42^. The three critical kinetochore linker proteins CENP-C^Mif2^, CENP-T^Cnn1^, and CENP-U^Ame1,^ are often lost or significantly diverged during evolution. In spite of the observed loss or significant divergence of the protein sequence, other molecular innovations of proteins to link centromeric chromatin to the outer kinetochore, ensuring accurate kinetochore-microtubule interactions remain largely unknown^22, 40, 43^.

Here, by analyzing a number of sequenced fungal genomes, we reveal that while retaining CENP-C^Mif2^, other CCAN components including conventional linker proteins CENP-T^Cnn1^ and CENP-U^Ame1^ homologs are lost multiple independent times in basidiomycetes. Furthermore, we identified a group of basidiomycete kinetochore proteins (Bkts) using *Cryptococcus neoformans*, a basidiomycete and human fungal pathogen, which undergoes a step-wise kinetochore assembly on a long repetitive regional centromere^44–47^ as a model system. Based on the further functional analysis performed *in vivo* and *in vitro,* we demonstrate that a novel protein bridgin^Bkt1^ connects the outer kinetochore to centromeric chromatin following its recruitment by KMN network proteins. Although being identified in a fungus, we predict the existence of bridgin homologs outside the fungal kingdom. These findings provide evidence of an unconventional kinetochore linker pathway, carried out by a novel class of kinetochore proteins found across eukaryotes.

## RESULTS

### Multiple independent loss events of CCAN proteins in Basidiomycota

To have a comprehensive understanding of the kinetochore composition in the fungal phyla of Basidiomycota, we analyzed putative kinetochore homologs using high confidence protein homology searches combined with secondary and tertiary structure prediction amongst species representing 31 fungal orders across the three sub-phyla (Pucciniomycotina, Ustilagomycotina, and Agaricomycotina). Additionally, representative species across 7 fungal phyla were considered. CENP-A^Cse4^, the 16-member CCAN, and the 10-member KMN network were chosen for this study (Fig 1a and Supplementary Table 1). Our analysis indicates robust conservation of the KMN network proteins across basidiomycetes, which is consistent with previous studies^39, 42^.

**Figure 1.**
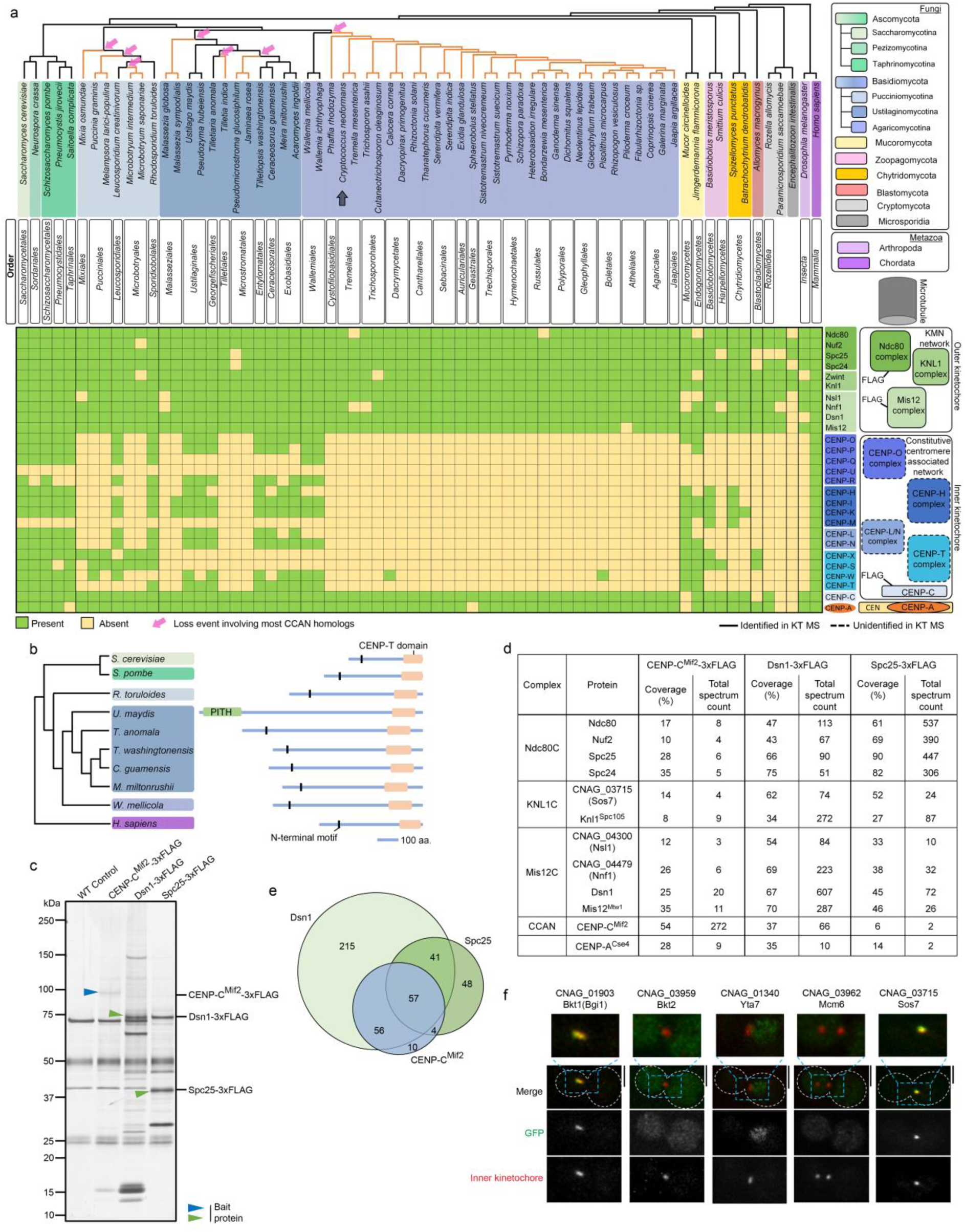
Identification of the kinetochore interactome in *C. neoformans*. **(a)** Conservation of kinetochore proteins across mentioned species. Cladogram representing the relationship between the species is drawn, and each sub-phylum is color-coded. Presence (green boxes) or absence (yellow boxes) of each kinetochore protein is shown. Pink arrows indicate loss events of CCAN proteins excluding CENP-C^Mif2^. FLAG labeling refers to proteins of kinetochore sub-complexes (CENP-C, Mis12C, and Ndc80C) tagged in *C. neoformans* and used for the IP-MS identification of the kinetochore interactome. Protein complexes identified (solid borders) or remained unidentified (dotted borders) by the IP-MS experiment in *C. neoformans* are shown. Grey arrow points to *C. neoformans.* Orange lines in the cladogram refer to basidiomycete lineages that have lost most CCAN components. **(b)** Domain architecture of identified CENP-T^Cnn1^ homologs amongst basidiomycetes. **(c)** A silver-stained gel shows kinetochore interacting proteins in the thiabendazole treated G2/M enriched cell population in *C. neoformans*. Proteins bound to each of the 3xFLAG tagged bait proteins were separated on a gradient PAGE gel. The left-most lane of the untagged control strain shows commonly found contaminating proteins in the single-step FLAG affinity purification elute. **(d)** List of kinetochore proteins with the percentage of amino acid sequence coverage and the number of total peptides specific to the corresponding protein obtained by MS analysis. **(e)** Venn diagram of identified interacting proteins. Circles in green indicate components of the KMN network, Dsn1, and Spc25, while members of the CCAN, CENP-C^Mif2^, are shown in a blue circle. **(f)** Micrographs of *C. neoformans* cells at metaphase expressing GFP tagged proteins identified by the screen mentioned above. Kinetochores are marked by a mCherry tagged inner kinetochore proteins, CENP-C^Mif2^ for Bgi1(Bkt1), Bkt2 and Yta7 or CENP-A^Cse4^ for Mcm6 and Sos7. Scale bar, 3 μm.

On the other hand, we observed that most CCAN proteins were recurrently lost across 23 basidiomycete orders (Fig 1a). In the sub-phylum of Agaricomycotina, which *C. neoformans* belongs to, the loss event may have occurred early at the time of divergence of Wallemiales from other orders. While in the sub-phyla of Pucciniomycotina and Ustilogamycotina, the loss of most CCAN subunits might have taken place on multiple independent occasions, as suggested by retention of these proteins in a few discrete orders (Fig 1a). The inner kinetochore linker protein CENP-C^Mif2^ was the only uniformly conserved CCAN component present across basidiomycetes. Other known linker proteins, CENP-T^Cnn1^ and CENP-U^Ame1,^ were often lost together. Although the primary protein sequence conservation is low among CENP-T^Cnn1^ homologs, they share a typical protein architecture, an N-terminal α-helix composed of conserved hydrophobic residues and the CENP-T^Cnn1^ motif at the C-terminal (Fig 1b). In the order Ustilaginales, a PITH domain spans the N-terminal of the CENP-T^Cnn1^ homolog consisting of an α-helix composed of conserved hydrophobic residues (Fig 1b). Thus, the CENP-C^Mif2^ linker pathway is the single known linker pathway among 23 of the 32 basidiomycete orders investigated. These observations suggest a recurrent loss of most CCAN proteins in Basidiomycota.

### Interactome of kinetochore components validates our prediction of the kinetochore composition in *C. neoformans*

We sought to validate our prediction of the kinetochore composition using a relatively well-studied basidiomycete yeast *C. neoformans.* In order to comprehensively determine the constitution of the kinetochore and its interactome *in vivo* we generated strains in which the endogenous genes of CENP-C^Mif2^, Dsn1 (Mis12C) and Spc25 (Ndc80C) were replaced with a C-terminus 3xFLAG tagged version and confirmed their functionality in the haploid *C. neoformans* type-strain background of H99α (Supplementary Fig. 1a-c). Mass spectrometry (MS) analyses was performed after FLAG immunoprecipitation (IP) of CENP-C^Mif2^, Dsn1, and Spc25 from metaphase enriched cell population, mitotic index >90% (Fig. 1c and Supplementary Table 2). All predicted inner kinetochore (CENP-A^Cse4^ and the CCAN component CENP-C^Mif2^) and outer kinetochore KMN network components were identified from each of the three FLAG purifications (Fig. 1d). Identified KMN network components included previously unannotated ORFs coding for proteins of the Mis12C (CNAG_04300^NSL1^, CNAG_04479^NNF1^) and the KNL1C (CNAG_03715^SOS7^) (Fig. 1d and Supplementary Table 2). Except for CENP-C^Mif2^, no components of the CCAN were identified (Fig. 1d, and Supplementary Table. 2). This result further validates that *C. neoformans* retains a single known kinetochore linker pathway through CENP-C^Mif2^ while having lost other known linker pathways, mediated by the CENP-T^Cnn1^ and CENP-U^Ame1^ complexes.

### A novel basidiomycete kinetochore protein (Bkt), Bridgin (Bgi1) is identified

We next investigated whether there existed unknown kinetochore proteins compensating for the loss of CCAN subunits in *C. neoformans.* Having identified all known structural kinetochore components from each IP-MS experiment, we hypothesized that it was possible to identify novel kinetochore proteins from the common list of interactors from CENP-C^Mif2^, Dsn1 and Spc25 affinity purification experiments (Fig. 1e). With this criteria we categorized two sets of proteins: a) primarily conserved amongst basidiomycetes with unknown function, and named them basidiomycete kinetochore proteins (Bkts) (CNAG_01903^Bkt1^, CNAG_03959^Bkt2^ and CNAG_02701^Bkt3^) (Supplementary Fig. 1d), and b) known chromatin interacting proteins with uncharacterized kinetochore function (CNAG_01340^Yta7^ and all components of the Mcm complex [Mcm2-7]) (Supplementary Fig. 1e). We used CNAG_03962^Mcm6^ as a representative test candidate. In the secondary screen, these five proteins were epitope-tagged at the native locus each with a C-terminus V5-GFP in a strain expressing mCherry-tagged inner kinetochore proteins, CENP-A^Cse4^ or CENP-C^Mif2^. CNAG_03715^SOS7^ was used as a candidate to validate the localization of identified unannotated kinetochore proteins (Fig. 1f and Supplementary Fig. 1f). Among the five proteins, strains where subcellular localization could be observed, was obtained for four, excluding CNAG_02701^Bkt3^. The protein encoded by the ORF.CNAG_01903^BKT1^ colocalized at specific cell cycle stages with the inner kinetochore protein, CENP-C^Mif2^ (Fig. 1f and Supplementary Fig. 1g). Other proteins did not show exclusive kinetochore localization, although some puncta of Yta7 and Mcm6 co-localized transiently with the inner kinetochore marker at G1/S stage of the cell cycle (Supplementary Fig. 1g). Based on the localization analysis of the protein hits, CNAG_01903^Bkt1^ was taken forward as a candidate kinetochore protein. Considering its identified function through this study, we refer to Bkt1 as bridgin (Bgi1) henceforth.

### Bridgin is recruited via multiple outer kinetochore receptors and its kinetochore level peaks at anaphase

Bridgin localized at the centromere in an M phase enriched cell population (Fig. 2a and Supplementary Fig. 1h). We subsequently investigated where bridgin localizes at the kinetochores in the kinetochore localization hierarchy. Our previous study suggested a step-wise assembly of the kinetochore, yet no sequential assembly of kinetochore sub-complexes was established in *C. neoformans*^44^. Using a microscopy-based interdependency analysis (Fig. 2b), we determined that the Mis12C and Ndc80C influence the stability of each other at the kinetochore (Supplementary Fig. 2b and c). The Dam1C (Supplementary Fig. 2h and i) and KNL1C (Supplementary Fig. 2d and e) independently require the Mis12C-Ndc80C platform for kinetochore recruitment. While the Ndc80C-Mic12C platform needed the inner kinetochore protein CENP-C^Mif2^ for its localization (Supplementary Fig. 2a). Performed interdependencies are tabulated (Supplementary Fig. 2k) and summarized in Supplementary Fig. 2l.

**Figure 2.**
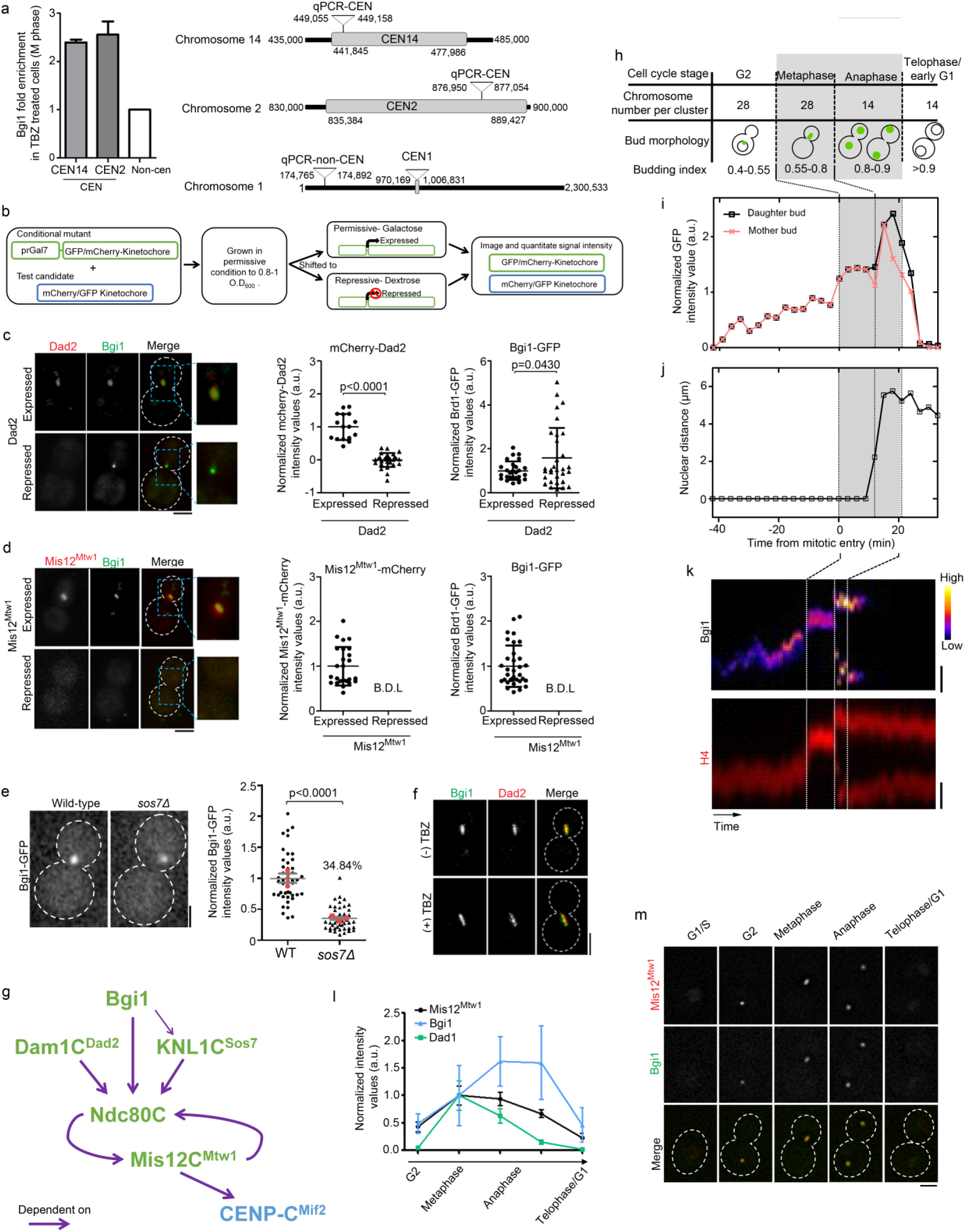
A novel kinetochore bridgin assembles onto the KMN network, reaching a maximum concentration during anaphase *C. neoformans*. **(a)** Measurement of levels of bridgin at the kinetochore in M phase cells by cross-linked ChIP-qPCR. Centromeres (CEN 14 and 2) and control non-centromeric regions were amplified by specific primers to determine its levels at the centromere. *N*=3. Error bars, s.d. Schematic is representing the location of centromeric-specific qPCR primers *(right)*. **(b)** Schematic of the experimental design to determine localization interdependency amongst kinetochore sub-complexes. **(c)** Bridgin localization signals in representative cells in the presence and absence of Dad2 (*left*). Levels of Dad2 and bridgin signals upon Dad2 expression and repression are quantified. For Dad2 measurements, *N*=16 and *N*=29 or Dad2 expressed and repressed conditions, respectively. For Bgi1-GFP measurements, *N*= 26 and *N*=31 for Dad2 expressed and repressed conditions respectively. *P*-value was determined using two-tailed t-test. Error bars, s.d. **(d)** Normalized intensities of Mis12^Mtw1^ and bridgin under conditionals of Mis12^Mtw1^ expression and repression. Strong influence of Mis12^Mtw1^ kinetochore protein on bridgin resulted in signals that were below detectable levels (B.D.L). Scale bar, 3 μm. *N*=25 and *N*=33 for Mis12^Mtw1^ and bridgin, respectively. Error bars, s.d. **(e)** Bridgin signal intensities at the kinetochore were measured in WT and *sos7Δ* cells. Red dots indicate the mean bridgin signal intensities of three independent transformants. Error bars, s.e.m. *P*-value was determined using two-tailed t-test. *N*=45. Scale bar, 2 μm. **(f)** Representative cells illustrate non-reliance of bridgin on an intact mitotic spindle for kinetochore localization. (+) TBZ cells were treated with 10 μg/ml thiabendazole (TBZ) for 3 h. **(g)** Schematic describes the observed localization interdependency of kinetochore protein complexes at the *C. neoformans* kinetochore. Protein complexes indicated in green or blue correspond to the outer or inner kinetochore, respectively. **(h)** Expected chromosome number per kinetochore cluster in the haploid type-strain H99α and spatial location of bridgin with corresponding budding index are tabled. **(i)** Signal intensity measurements of bridgin from G2 phase until the subsequent G1 phase are shown in a plot at an interval of 3 min. The event of mitotic entry is referred to as time point 0. **(j)** A cell is considered to have exited anaphase when nuclear distances have reached their maxima. **(k)** A kymograph of the corresponding tabulated bridgin signals. Time interval represented, 1 min for a total of 100 min. Scale bar, 2 μm. **(l)** Comparison of protein levels of bridgin, and representative proteins of the outer kinetochore. *N*=5 each kinetochore protein. Error bars, s.d. **(m)** Co-localization of Bgi1-GFP signals in cells at the various stages of the cell cycle with Mis12^Mtw1^ in an asynchronous culture. Scale bar, 3 μm.

Subsequent interdependency analyses suggested that bridgin localizes to the kinetochore independent of Dad2 (Dam1C) (Fig. 2c). On the other hand, bridgin localization at the kinetochore wholly and partially (∼65%) depends on Mis12^Mtw1^(Mis12C) (Fig. 2d) and Sos7(KNL1C) (Fig. 2e), respectively. Further, bridgin’s kinetochore localization is independent of spindle integrity (Fig. 2f). The recruitment of bridgin downstream of the KNL1C and Mis12C-Ndc80C platform suggests that there may exist multiple binding sites for bridgin onto the KMN network (Fig. 2g).

To understand how bridgin dynamics is regulated during cell cycle progression, we analyzed bridgin signal intensities at the kinetochore. Bridgin localized to the kinetochore starting from G2 until telophase/G1 (Fig. 2h-k and Supplementary Fig. 1j). The signal intensities of bridgin at the kinetochore reached a peak immediately at the onset of anaphase, attaining an average ∼150% of metaphase intensities (Fig. 2i, k and l, and Supplementary Fig.1j). The dynamic intensities of other transiently localized kinetochore proteins of the KMN network, Mis12^Mtw1^, Nuf2, and Knl1^Spc105^, and the subunits of the Dam1C, Dad1, and Dad2, were measured. The KMN network proteins localized concomitantly to the kinetochore during G2 and persisted until telophase/G1, reaching the maximum signal intensity during metaphase (Fig. 2l and Supplementary Fig. 1i). The Dam1C proteins Dad1 and Dad2 localized post-mitotic onset exclusively, reaching peak intensities at metaphase and reducing sharply, almost to an undetectable level in late anaphase (Fig. 2l and Supplementary Fig. 1i). Analysis of bridgin localization in an asynchronous population further validated the cell cycle-stage specific kinetochore localization, similar to the outer kinetochore protein Mis12^Mtw1^ (Fig. 2m). Yet, KMN network proteins and bridgin reached peak kinetochore intensities at distinct times, metaphase and anaphase, respectively (Fig. 2l).

Enrichment at the kinetochore, dependence on outer kinetochore KMN network proteins and spindle independent kinetochore localization of bridgin strongly implicate that bridgin is an outer kinetochore protein. Further, taking into consideration the interdependency with the outer kinetochore proteins and the localization dynamics, we propose that bridgin localizes onto the KMN platform at the kinetochore.

### Bridgin is essential for accurate kinetochore-microtubule interaction

Bridgin-null (*bgi1Δ*) strains were generated to characterize the function of bridgin as a kinetochore protein. *bgi1Δ* cells exhibited reduced growth rates (Fig. 3a), and ∼20% loss in viability (Supplementary Fig. 3a). These mutant cells also displayed a ∼90-fold increase in the gross missegregation rate, which may account for the reduced viability in *bgi1Δ* (Fig. 3b and Supplementary Fig. 3b). *bgi1Δ* defects were complemented by the reintegration of the full-length bridgin gene (Bgi1^FL^) expressed under its native promoter (Fig. 3b and Supplementary Fig. 3a). We subsequently examined how *bgi1Δ* affected cell cycle progression. For this, microscopic markers to determine cell cycle stages previously determined^44, 48^ were used and summarized in Supplementary Fig. 3c. While WT cells spent an average of 18 min in M phase, *bgi1Δ* cells showed a delay in M phase for ∼30min (an under-representation since ∼10% of cells failed to exit M phase arrest even after >50 min) (Supplementary Fig. 3d). Within the M phase, the delay occurred prior to anaphase onset (Fig. 3c and Supplementary Fig. 4f). *bgi1Δ*, when examined by live-cell microscopy, exhibited unattached chromosomes, lagging chromosomes, and micronuclei formation defects, suggesting that inaccurate kinetochore-microtubule attachments occurred (Fig. 3d-f and Supplementary Fig. 3f). *mad2Δ* in the background of *bgi1Δ* alleviated M phase delay (Fig. 3c and Supplementary Fig. 3f), but the double mutants were conditionally synthetic lethal upon treatment of the microtubule poison thiabendazole (2 μg/ml) or under conditions of spindle insult (14°C and 37°C) (Fig. 3g). Based on these observations, we conclude that bridgin is essential for accurate kinetochore-microtubule interactions. Absence of bridgin in cells (*bgi1Δ*) elicit a prolonged SAC response in its absence, to correct for erroneously kinetochore-microtubule attachments.

**Figure 3.**
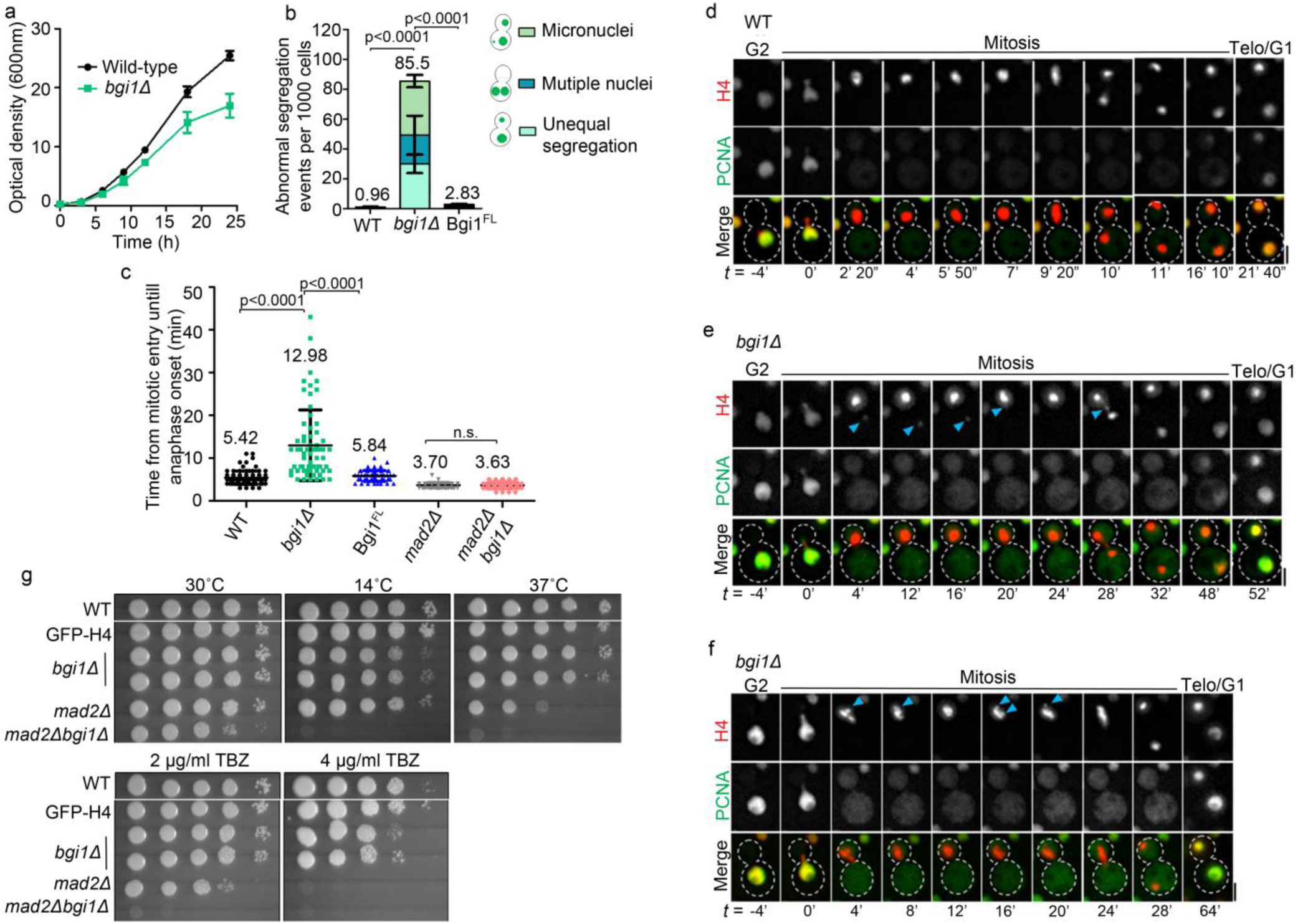
Loss of bridgin impairs kinetochore-microtubule interactions. **(a)** Growth curve of the WT strain H99α and bridgin null mutant (*bgi1Δ*) cells at 30°C. *N*=3. **(b)** The rate of abnormal nuclear segregation events was measured at 30°C using GFP-H4 in WT, *bgi1Δ,* and bridgin full-length re-integrant (Bgi1^FL^) strains. The number of cells examined was >3000, *N*=3. Error bars, s.d. *P*-value was determined using two-tailed t-test. **(c)** Time from the mitotic entry to anaphase onset was quantified and plotted for each strain as indicated. Diffusion of a nuclear marker PCNA coincided with the migration of the nucleus into the daughter cell, indicating entry into semi-open mitosis. Nuclear distance >1 μm was considered as the entry into anaphase. Mean of WT (*N*=88), *bgi1Δ* (*N*=64), Bgi1^FL^ (*N*=61), *mad2Δ* (*N*=67) and *mad2Δ bgi1Δ* (*N*=56) were measured as indicated. Error bars, s.d. *P*-value was determined using two-tailed t-test. Not significant, n.s. **(d-f)** Representative time-lapse images of **(d)** WT or **(e and f)** *bgi1Δ* cells. Time measurements were made from the mitotic onset. Cell cycle stages were scored for either by PCNA localization or chromatin condensation (H4-GFP) and nuclear migration into the daughter bud. Scale bar, 2 μm. **(e)** Blue arrows point to an unattached chromosome at the mitotic onset and a lagging chromosome at anaphase. **(f)** Blue arrows indicate a chromosome that is separated from the compact chromatin mass in prometaphase as seen in WT. **(g)** Serial 10-fold dilutions starting from 2×10^5^ cells were spotted for WT, GFP-H4, *bgi1Δ*, *mad2Δ*, and *mad2Δ bgi1Δ* are shown.

### Basic domain of bridgin is dispensable for kinetochore localization but critical for its function

We next sought to understand how bridgin carries out its role at the kinetochore. Bridgin has 1295-amino acid (aa), in which aa1-124 forms a fork-head associated (FHA) domain (a phosphopeptide recognition domain), aa161-164 creates an unconventional putative PP1 docking site, while the remaining region of the protein is predicted to be largely unstructured (Fig. 4a). The aa1005-1295 C-terminal region is predicted to have a pI of 11.20, and we refer to this region as the basic domain (BD). The unstructured domain (USD) was defined as the region spanning aa125-1004, which was acidic with a pI of 4.65 (Fig. 4a) and contained 13 repeats of a bridgin consensus motif (Supplementary Fig. 4a and Supplementary Table 3). Domain deletion constructs were generated as described in Supplementary Fig. 4b, wherein the domain deletion is expressed under its native promoter with an N-terminus 3xFLAG-GFP epitope tag. The cassettes were reintegrated into *bgi1Δ* cells expressing H4-mCherry (Supplementary Fig. 4b) to obtain strains expressing truncated bridgin proteins lacking various domains, as mentioned in Fig. 4b. Microscopic estimation of GFP signal intensities of the bridgin derivatives and the Bgi1^FL^ suggested that the FHA domain (FD) and USD regions were able to localize independently of each other at the kinetochore, albeit to different extents of ∼20% and ∼40% of the WT level respectively (Fig. 4c and d). Localization of Bgi1^BDΔ^ at the kinetochore was not significantly different from that of WT (Fig. 4c and d). Further, lack of kinetochore localization by the BD suggested it was not involved in kinetochore localization of bridgin. Localization analyses using various truncated mutants suggest that bridgin through its FD and USD can make multiple contacts at the kinetochore, consistent with the observation that bridgin is recruited downstream to multiple outer kinetochore KMN proteins (Fig. 2g).

**Figure 4.**
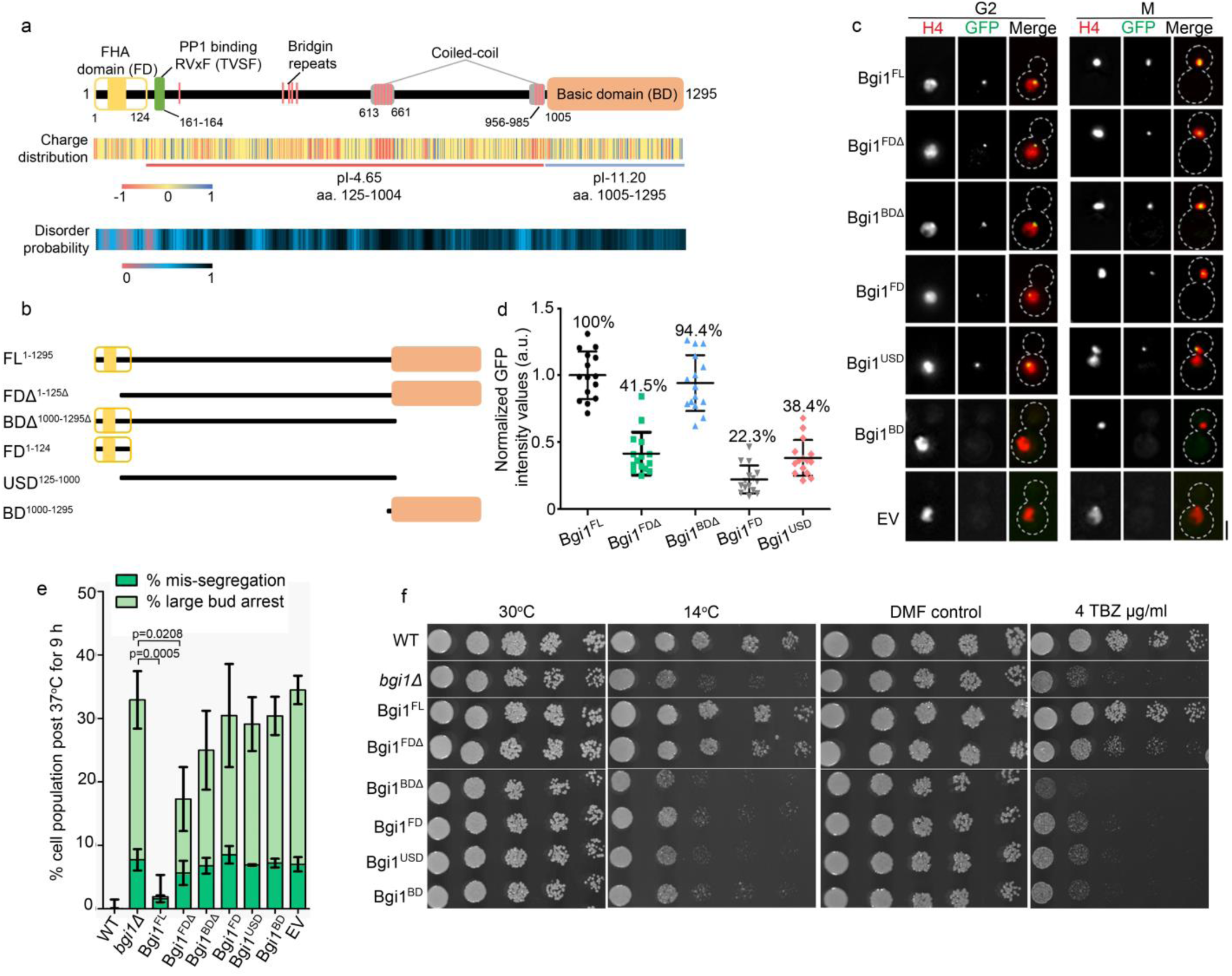
Kinetochore recruitment and positioning of bridgin via multiple receptor sites are essential but insufficient for its function. **(a)** Schematic describing predicted features of bridgin protein. Charge distribution of amino acid residues was predicted with a window size of 2 using EMBOSS charge. Disorder probability of bridgin was calculated using IUPRED2A. **(b)** Schematic of generated domain deletion constructs of bridgin. Constructs were generated with a 3xFLAG-GFP tag at the amino-terminus. **(c)** Representative cells in G2 and M phase showing localization of the mentioned bridgin or its derivatives and the control empty vector (EV). Nuclear localization was scored for using the chromatin marker histone H4. Scale bar, 3 μm. **(d)** Bridgin signal intensities were measured in 15 mitotic cells prior to anaphase onset. Percent intensity values normalized to Bgi1^FL^ have been mentioned. **(e)** The extent of complementation by truncated proteins of bridgin was measured. H4-mCherry was localized in bridgin protein derivatives, lacking various domains, in a *bgi1Δ* strain background. Cells were grown at 30°C until log phase and transferred to 37°C. Indicated cell populations were measured 9 h post-incubation at 37°C. All values were normalized to WT. Defects in nuclear segregation were measured as mentioned in Fig 3b. Error bars, s.d. The number of cells examined was >1000, *N*=3 for each indicated strain. *P*-value was determined using two-tailed t-test. **(f)** Cells of varying numbers 2×10^4^, 2×10^3^, 200, 100 and 50 cells were spotted on plates as indicated.

Subsequently, to define the domains necessary for bridgin function, we scored for complementation of the *bgi1Δ* phenotype at 37°C (used for ease of scoring due to enhancement in the population of M phase delayed cells) (Fig. 4e) and cell growth assays under conditions altering microtubule dynamics (Fig. 4f). Partial complementation of phenotype was observed for the Bgi1^FDΔ^ mutant, exhibiting reduced kinetochore localization while retaining the BD. No significant complementation was obtained for any of the other domain deletion constructs including the Bgi1^BDΔ^, that localized to the kinetochore similar to Bgi1^FL^ levels (Fig. 4e and f). Bgi1^FL^ was able to suppress *bgi1Δ* phenotype significantly (Fig. 4e and f). Comparable results were observed for the rate of missegregation events at 30°C (Supplementary Fig. 4c), albeit weak complementation was observed for Bgi1^BDΔ^. Taken together, all domains, including BD, which is not related to kinetochore localization of bridgin, are critical for the full function of bridgin.

### The C-terminal basic domain of bridgin interacts with DNA

Impact of bridgin on SAC activity (Supplementary Fig. 4d and e), spindle dynamics (measured at its most dynamic stage, anaphase, Supplementary Fig. 4f) and gross kinetochore composition (Supplementary Fig. 4g-j) were tested and found to be unaltered in *bgi1Δ.* Thus these factors were ruled out as possible reasons of defective kinetochore-microtubule attachments associated with *bgi1Δ* mutants.

To address the role of BD towards bridgin function, FLAG affinity purification of Bgi1^FL^ (using 150 mM KCl and a more stringent condition of 300 mM KCl) and Bgi1^BDΔ^ (150 mM KCl) was performed and these samples were subject to mass spectrometry analysis (Fig. 5a, Supplementary Fig. 4k and Supplementary Table 4). A comparison of the relative abundance of specific interactors obtained within Bgi1^FL^ and Bgi1^BDΔ^ suggested an enrichment of chromatin interacting proteins in Bgi1^FL^ affinity purification over Bgi1^BDΔ^ (Fig. 5b *top*). While, kinetochore proteins were relatively more abundant as interactors in Bgi1^BDΔ^ over Bgi1^FL^ affinity purifications (Fig. 5b *bottom*), with proteins of the KMN network being the top hits (Fig. 5b and Supplementary Table 4).

**Figure 5.**
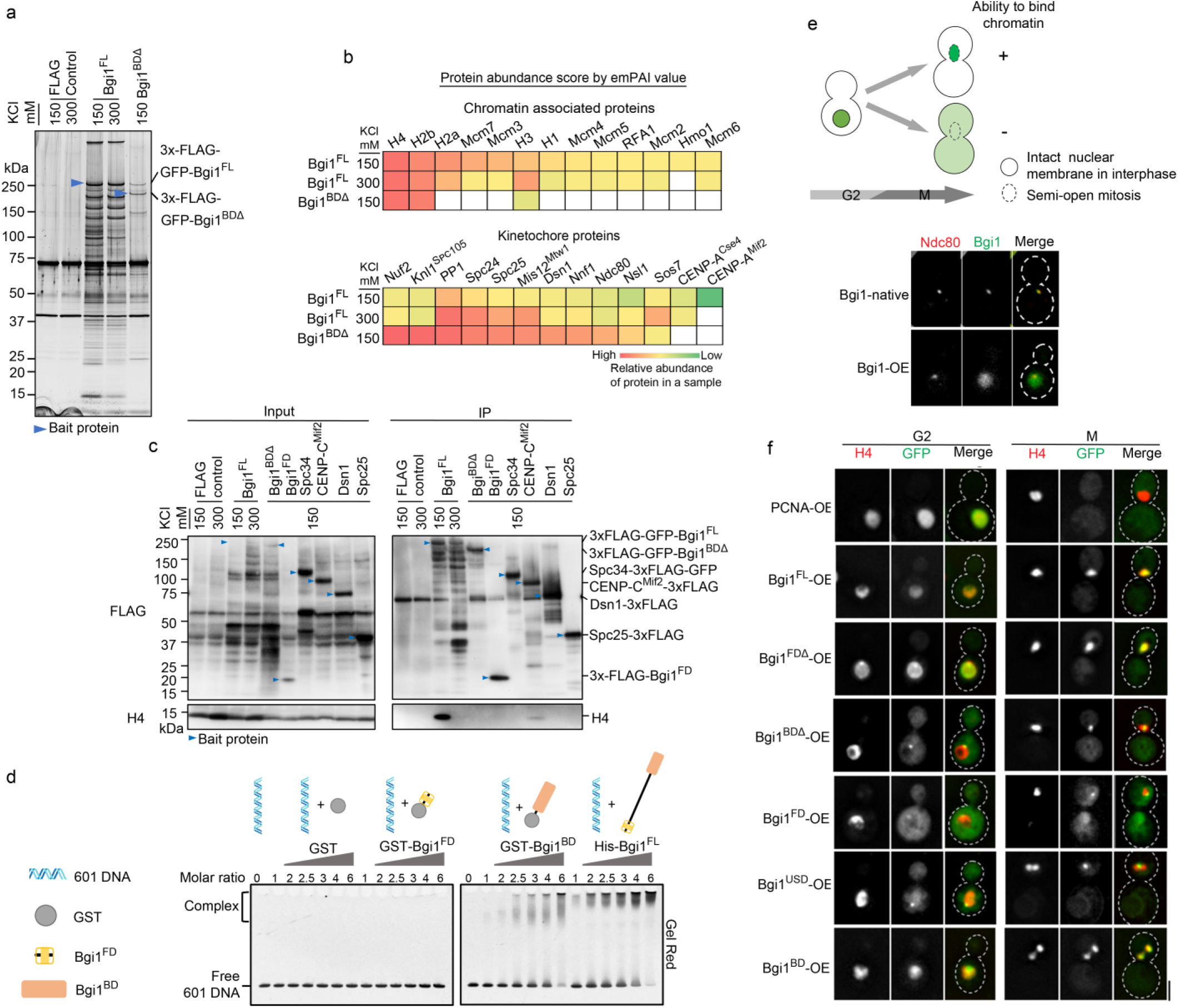
The basic carboxy terminus of bridgin has a property to interact with DNA non-specifically *in vitro* and *in vivo.* **(a)** The silver-stained gel used to visualize bridgin interacting proteins. Blue arrows indicate the bait protein on each lane as applicable. Lysates for the immunoprecipitation experiment were prepared from a G2/M cell population that was enriched by treatment with 10 μg/ml of TBZ for 3 h. Two left lanes show common contaminating proteins obtained in the single-step 3xFLAG affinity purification. **(b)** List of chromatin-associated and kinetochore proteins obtained as interactors from bridgin affinity purification. Top 10 known chromatin-associated proteins obtained in Bgi1^FL^ 150 mM affinity purification *(top)* and known kinetochore proteins obtained in Bgi1^BDΔ^ 150 mM affinity purification were chosen and arranged in ascending order from left to right based on relative abundance scores across each IP. The scores are based on emPAI values obtained for interacting proteins across affinity purifications. Undetected proteins are represented by white boxes. **(c)** Proteins from 3xFLAG tagged strains were extracted, and affinity purifications were performed with FLAG antibodies. Interacting proteins were eluted with 3xFLAG peptides, and blots were probed with FLAG and H4 antibodies. Bait protein bands are indicated. **(d)** Electrophoretic mobility shift assays (EMSA) samples were separated on a PAGE gel and stained with Gel Red for visualization. **(e)** Chromatin-bound proteins colocalize with the nuclear marker H4-mCherry in metaphase while free nuclear proteins diffuse into the cytoplasm following the entry into mitosis *(top)*. Visualization of bridgin localization by fluorescence microscopy when expressed under the native or an Over-expression (OE) promoter construct. Outer kinetochore protein Ndc80 was used to mark the kinetochore. **(f)** Visualization of Bgi1-OE constructs. GFP-Bgi1-OE constructs were transformed into the H4-mCherry *bgi1Δ* strain. Representative images of cells in G2 and M phase are shown. Scale bar, 3 μm.

Based on the observation that chromatin interacting proteins are more enriched in the Bgi1 construct containing the BD, Bgi1^FL^, we hypothesized that BD might interact with chromatin. Through co-immunoprecipitation experiments, histone H4 was found to associate with Bgi1^FL^ (150 mM) and to a reduced extent with CENP-C^Mif2^ (Fig. 5c). No detectable association of histone H4 was obtained with outer kinetochore proteins (Dsn1, Spc25, and Spc34) or Bgi1^BDΔ^ (Fig. 5c). These results led us to hypothesize that bridgin-chromatin interaction occurs through Bgi1-BD and not a consequence of bridgin receptor assembly, KMN network, onto centromeric chromatin. We tested the possibility of interaction between bridgin BD with chromatin *in vitro* by EMSA and found that the BD binds to DNA (Fig. 5d) and nucleosomes of varying compositions (Supplementary Fig. 5a). Further, these observations indicated to us that the interaction between the BD and DNA/chromatin might be non-specific.

While we observed non-specific bridgin-chromatin interactions *in vitro*, we hypothesized that if additional regulators existed *in vivo* to restrict bridgin BD localization to centromeric chromatin, over-expression (OE) of bridgin would not alter its localization (Supplementary Fig. 5b). On the contrary, we observed that localization of Bgi1-OE was transformed and overlapped with chromatin marked by histone H4 (Fig. 5e and Supplementary Fig 5c), indicating that bridgin can interact with DNA/chromatin non-specifically *in vivo* as well. Microtubule-like signal or localization of Bgi1-OE outside chromatin, marked by H4, was not observed. Hence, we ruled out the possibility of the outer kinetochore protein bridgin in binding to microtubules. Further, we used the over-expression strategy as an assay to determine the DNA binding ability of bridgin domain deletion mutants *in vivo* (Fig. 5f). PCNA was used as a negative control, wherein nuclear-localized but chromatin unbound PCNA pool diffused into the cytoplasm during M phase, on account of semi-open mitosis. While Bgi1^FL^-OE localization was observed to overlap with H4-mCherry, Bgi1^BDΔ^-OE was restricted to a punctum. Supporting the notion that bridgin localizes to the kinetochore through FD and USD, a punctum for both constructs, Bgi1^FD^ and Bgi1^USD^, were observed. Further, the localization of Bgi1^BD^ was found to be similar to Bgi1^FL^. Thus, these observations suggested that the BD was necessary and adequate to bind chromatin *in vivo,* and the loss of BD in the over-expression constructs was sufficient to restrict bridgin localization to the kinetochore puncta.

Considering bridgin was recruited to the outer kinetochore downstream of the KMN network, it was surprising that bridgin BD binds to chromatin. Further, increased enrichment of DNA from the Bgi1^FL^ over Bgi1^BDΔ^ in the native-ChIP suggested that bridgin, through its BD interacts with DNA when kinetochore localized (Supplementary Fig. 5d).

### Basic nature of bridgin BD is vital for its function

We show that bridgin loss does not alter previously described chromatin marks of H3K9me2 and CpG methylation at *C. neoformans* centromeres (Supplementary Fig. 5 e and f), towards understanding the consequence of bridgin binding to DNA. To summarize our findings, we observe that bridgin localizes to the kinetochore through FD and USD, and its interaction with DNA/chromatin through its BD is essential for its function (Supplementary Fig. 6a). However, it is still unclear how BD influences bridgin’s function. We hypothesize two possibilities: a) interaction of BD with other proteins at chromatin is essential for bridgin function or b) the ability of BD to interact with DNA is adequate for bridgin’s function. To distinguish these possibilities, we performed a domain-swap experiment. Replacing the BD^1005–1295^ of bridgin with an amino acid stretch of similar properties (length: ∼300aa., unstructured, non-specific DNA binding and a charge of ∼pI of 10) found in the basic region (BD)^2937–, 3256^ of the human Ki67 gene (Fig. 6a). Ki67 was previously shown to bind non-specifically to DNA^49^ and functions as a surfactant by coating chromosomes during mitosis^50^. We confirmed that Ki67 BD^2937–3256^ binds to DNA non-specifically in *C. neoformans* using the overexpression assay (Supplementary Fig. 6b). Bgi1^FL^, Bgi1^BDΔ^, and Bgi1^BDΔ^+Ki67^BD^ were expressed under the native bridgin promoter as described for other domain deletion constructs and found to localize to the kinetochore with similar intensities when integrated into a *bgi1Δ* background strain (Fig. 6b and c). Weak complementation was observed for Bgi1^BDΔ^ over *bgi1Δ* (Fig. 6d). On the other hand, the Bgi1^BDΔ^+Ki67^BD^ construct was able to complement defects observed in *bgi1Δ* and the Bgi1^BDΔ^ mutants. The Bgi1^BDΔ^+Ki67^BD^ phenotype was non-significant from the FL (Fig. 6d and Supplementary Fig. 6c). These observations were additionally validated by the spotting growth assay (Fig. 6e).

**Figure 6.**
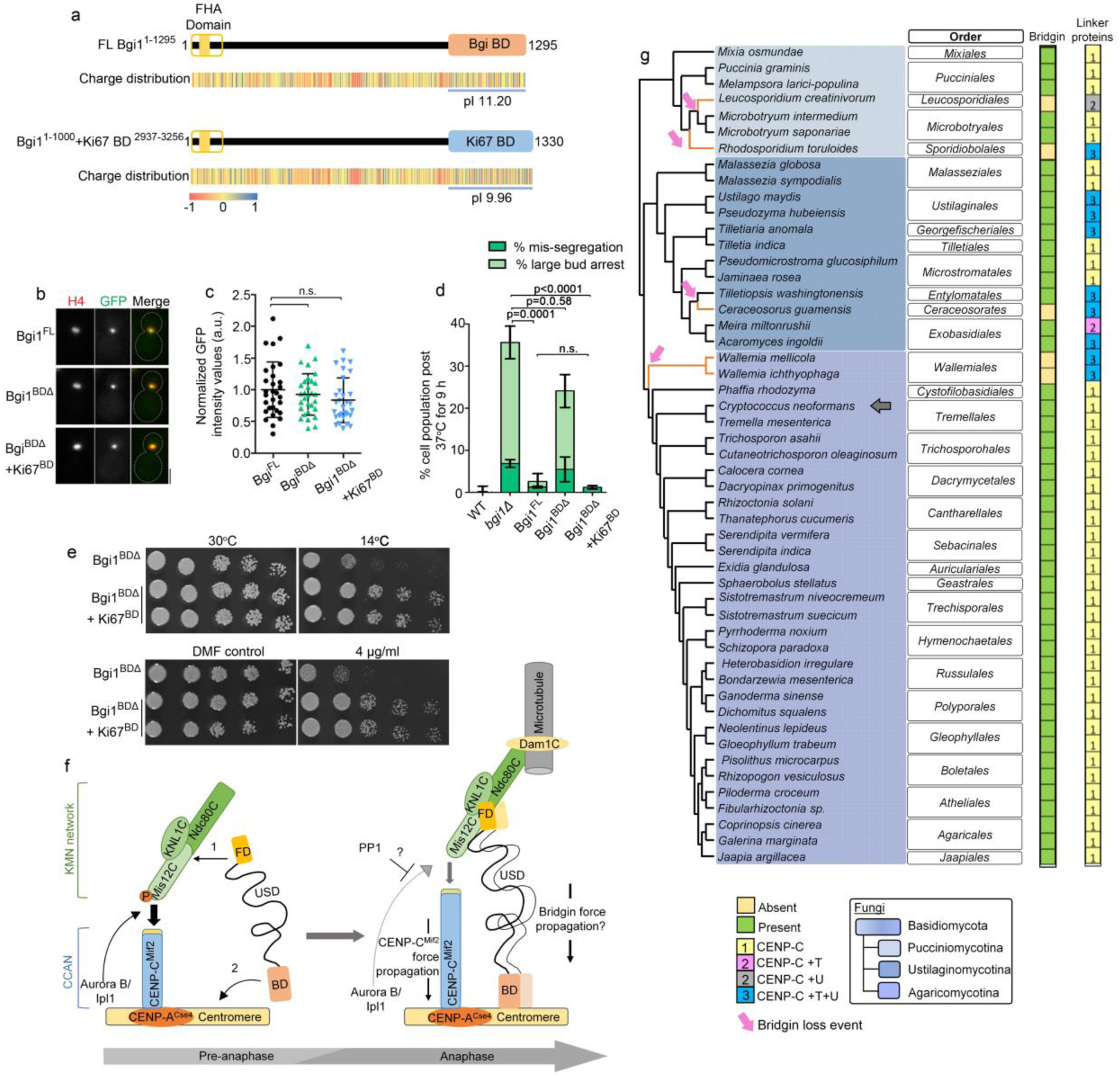
Bridgin is an unconventional linker of the outer kinetochore and centromeric chromatin. **(a)** Schematic representation of bridgin in which its basic domain aa1005-1295 with a pI of 11.2, was replaced with the basic DNA binding domain from HsKi67, aa2937-3256 that exhibits a pI of 9.96. **(b)** Representative micrographs of Bgi1^FL^, Bgi1^BDΔ^ and Bgi1^BDΔ^+Ki67^BD^, expressed in *bgi1Δ* cells expressing H4-mCherry. **(c)** Quantitation of Bgi1 and chimeric Bgi1 signals in 30 cells. Not significant (n.s.). *P*-value was determined using two-tailed t-test. Scale bar, 3 μm. **(d)** Complementation of *bgi1Δ* phenotype by Bgi1^FL^, Bgi1^BDΔ^ and Bgi1^BDΔ^+Ki67^BD^ protein derivatives was measured by assessing their phenotype post-incubation of cells to 37°C for 9 h. Error bars, s.d. The number of cells examined was >1000, *N*=3 for each indicated strain. *P*-value was determined using two-tailed t-test. Not significant (n.s.) **(e)** Cells of varying numbers 2×10^4^, 2×10^3^, 200, 100 and 50 were spotted on YPD without TBZ and YPD containing 4 μg/ml TBZ. **(f)** A model describing bridgin as a kinetochore protein connecting the outer KMN network, through its FD and USD, and directly to DNA via its basic DNA binding domain. Restricted interaction of bridgin with DNA in WT cells is a possible consequence of outer kinetochore specific recruitment prior to its interaction with centromeric chromatin. Increased bridgin localization is observed in anaphase **(g)** Identification of bridgin homologs across Basidiomycota. Presence or absence of a bridgin homologs is represented. No of identified linker pathways are mentioned and color-coded to represent the linker pathway(s) present. Grey arrow points to *C. neoformans.* Bridgin loss events in basidiomycete lineages are represented by orange lines in the cladogram.

While we cannot rule entirely out the contribution of the amino and/or middle region of bridgin towards function, independent of its kinetochore localization capacity. We propose that bridgin ensures accurate kinetochore-microtubule attachments and mitotic fidelity by linking the outer kinetochore to centromeric chromatin, based on the ability of bridgin to simultaneously localize to the outer kinetochore through the KMN network and to bind to DNA (Fig. 6f).

## DISCUSSION

As a step towards understanding the evolution of kinetochore organization and composition, we chose to study the kinetochore interactome of the human pathogen and a basidiomycete yeast *C. neoformans.* During the study, we identified a novel outer kinetochore protein, that we termed as bridgin in *C. neoformans*. Our experiments strongly indicate the absence of most known CCAN proteins, with the exception of CENP-C^Mif2^, and presence of all KMN network proteins in this system, validating our bioinformatic prediction. Thus, suggesting a single known linker pathway (CENP-C) from centromeric chromatin to the outer kinetochore, reminiscent of the fruit fly *D. melanogaster* and the nematode *C. elegans* like kinetochores^39, 51–54^. However, we propose, and as the name suggests, that bridgin functions as a new linker protein, since it binds to the outer kinetochore and centromeric DNA simultaneously, analogous to previously described linker proteins CENP-C^Mif2 33, 55, 56^ and CENP-T^Cnn1 15, 40, 57^. Presence of multiple kinetochore linker pathways is critical, to varying extents, in overcoming Dsn1 inhibition^29, 30, 32^. Unlike the single linker pathway containing kinetochore such as that of *D. melanogaster*^52, 58^, *C. neoformans* retains the Dsn1 autoinhibitory domain (Fig. 1a and Supplementary Fig. 6d). Although a recent study suggests Nnf1 to be the Dsn1 homolog is *D. melanogaster*, we were unable to identify the presence of the Dsn1 autoinhibitory domain in the suggested homolog^39^. Through our findings, we propose a role for bridgin towards linking the outer kinetochore by its recruitment and interaction with multiple KMN network proteins and DNA thereby promoting accurate kinetochore-microtubule attachments in *C. neoformans* (Fig. 6f).

An inability of the BD to localize specifically to the kinetochore and non-reliance of bridgin on sequence specificity for BD function endorses the hypothesis that binding of bridgin BD to DNA is a consequence of specific kinetochore recruitment (Fig. 6f). Rather unique to bridgin as a linker protein is the fact that its kinetochore localization is dependent on conserved KMN network proteins, Sos7 (KNL1C), and the Mis12C-Ndc80C platform (Fig. 2d, e, and g). CENP-T homologs require other CCAN proteins for its kinetochore localization^29, 59, 60^ and binds non-specifically to DNA *in vitro.* CENP-T^Cnn1^ was shown to increase the stability of a mini-chromosome possibly due to its ability to recruit the Ndc80C as suggested in a recent study ^29, 40^, while CENP-T in metazoans was shown to recruit the KMN network when ectopically tethered^30, 32, 61^. Bridgin does not appear to influence the recruitment of outer kinetochore proteins (Supplementary Fig4. g-i), further supported by the lack of Ndc80 mislocalization upon recruitment of bridgin to ectopic sites (Fig. 5e and Supplementary Fig. 5c). Bridgin levels at the kinetochore reach a peak at anaphase (Fig. 2i and l), a time when Aurora B^Ipl1^-mediated phosphorylation is suggested to be countered by phosphatase activity (Fig. 6f). The sharp reduction of AuroraB^Ipl1^ localization at anaphase kinetochores^48^ and essentiality of amino-terminus of CENP-C^Mif2^ (Supplementary Fig. 6e and data not shown) are observed in *C. neoformans.* Taken together, we propose that the kinetochore architecture alters during the metaphase-anaphase transition and the bridgin linker pathway functions to reinforce/stabilize the outer kinetochore. Thus, an important question we must address in future is whether the presence of Dsn1 autoinhibition can provide a constraint driving evolution/maintenance of multiple outer kinetochore linker pathways required for outer kinetochore reinforcement in organisms with monocentric chromosomes.

Outer kinetochore proteins are found to be more conserved than their inner kinetochore counterparts, including linker proteins, across eukaryotes^39, 42^ (Fig. 1a). Thus, additional KMN recruited linker pathways like the bridgin-pathway may provide cells with an effective alternative towards outer kinetochore reinforcement. Bridgin homologs are identified across all basidiomycete sub-phylum (Fig. 6f and Supplementary Table 5). Strikingly, an inability to identify bridgin homologs is specific orders correlates with the presence of multiple known linker pathways (Fig. 6g). It would be worth investigating whether the presence of multiple linker pathways may have allowed for flexibility in the retention of specific linker pathways in basidiomycetes. Genome sequencing of a greater number of distinct basidiomycetes would help address the correlation. It would be intriguing to recognize the contribution of the multiple linker pathways in organisms like *U. maydis* which retained CENP-T, CENP-C in addition to bridgin.

The identification of bridgin homologs in the basal ascomycetes of the class Pneumocystidales, such as in *Pneumocystis jirovecii* (causative organism of pneumonia), and Taphrinales, and further, identification of bridgin-like proteins outside fungi may suggest a more ancient origin of this class of kinetochore proteins containing a FHA domain (Supplementary Fig. 6f). In metazoans, the identified proteins with bridgin-like architecture (an amino-terminal FHA domain followed by a PP1 docking site, an unstructured central region containing repeats and a basic carboxy-terminus, Supplementary Fig. 6f) was found to code for Ki67, a component of the mitotic chromosome periphery^50, 62^. The role of bridgin-like proteins outside Metazoa is not known.

Future experiments will have to reveal how bridgin’s kinetochore recruitment and DNA/chromatin binding is regulated and the extent of its ability to bear load at the kinetochore. Furthermore, towards understanding bridgin biology, addressing if bridgin has additional functions at the kinetochore, for example, through PP1 recruitment is essential. We also are looking forward to screening other hits obtained as part of the *C. neoformans* kinetochore interactome. We hope further studies on non-conventional model systems like *C. neoformans* will help reveal conserved fundamental principles of the kinetochore architecture and its organization.

## ONLINE METHODS

### Homolog detection

All searches were carried out in the NCBI non-redundant protein database or the UniProtKB. Searches for kinetochore homologs were initially carried out using iterative HMMER^63^ jackhammer searches (E-value ≤ 10^-3^) with Pfam models for the mentioned kinetochore proteins. When available models of both yeast and metazoan kinetochore homologs were considered. Obtained hits were validated by performing reciprocal HMMER searches. Secondary structure of obtained hits was validated using Jpred4 and tertiary structure prediction using HHpred^64^ and/or Phyre2^65^. Protein sequences which were unable to produce hits upon reciprocal searches or failed to conform to expected secondary and tertiary structures were discarded. Further searches were performed with the same criteria using identified homologs phylogenetically closest to the species in question. Species considered in the study are mentioned in Supplementary Table 1, when homologs were not identified from a specific strain, an obtained homolog from another strain of the same species was considered. If multiple splice variants were identified the longest splice variant was mentioned. Obtained hits when possible was validated with the identified homologs from *C. neoformans.* Known kinetochore homologs from *S. cerevisiae, S. pombe, D. melanogaster,* and *H. sapiens* were used to draw the matrix of kinetochore homologs.

Towards identifying homologs of bridgin, the conserved FHA domain was taken as the bait for subsequent iterative HMMER jackhammer searches. Obtained hits were further screened for overall protein architecture (amino-terminus FHA domain, an unstructured central region and a basic carboxy-terminus, Supplementary Fig. 6f). Probability of protein disorder was predicted using IUPred2A^66^ and pI of the amino acid residues was predicted using ProtParam^67^. Amongst pucciniomycetes, *Exidia glandulosa* and *Sistotremastrum suecicum* the basic C-terminus is ∼150 aa. in contrast to ∼300 in other organisms.

Using published multi-gene and genome-scale phylogenetic data from The Fungal Kingdom^68^, JGI MycoCosm^69^, Interactive Tree of Life (iTOL) v4^70^ and Wang, Q. M. *et al* ^71^ the cladograms were drawn showing the relationship amongst the considered species.

### Yeast strains and plasmids

A list of strains and plasmids used in the study can be found in Supplementary Table 6. Primers used to generate the constructs are mentioned in Supplementary Table 7.

Conditional kinetochore mutant strains were grown on 1 % yeast extract, 2 % peptone, and 2 % galactose (YPG). All other strains were grown in 1 % yeast extract, 2 % peptone, and 2 % dextrose (YPD) at 30°C, 180 rpm unless mentioned otherwise. Strains were maintained on YPD/YPG solidified with 2% agar and stored at 4°C or −80°C in 15% glycerol. Yeast strains are based on the haploid type strain H99α or KN99a and generated by the standard procedure as previously described^44^. In brief, created native tagging and Gal7 promoter replacement cassettes were excised from the plasmid construct, over-expression cassettes were linearized by appropriate restriction enzymes, and deletion cassettes were generated by overlap PCR and transformed into *C. neoformans* strains of appropriate background by biolistic transformation^72^. Transformed cells were selected on drug selection in YPG for Gal7 promoter^73^ replacement strains to generate conditional mutants and YPD for all other strains.

### Protein affinity purification and native chromatin immunoprecipitation

An overnight culture was inoculated at 0.1 OD_600_ into fresh YPD. Grown until ∼0.7 OD_600_ and treated with 10 μg/ml of thiabendazole (TBZ) for 3 h. Cells were harvest, washed once in water followed by one wash with binding buffer BB150 (25 mM HEPES pH 8.0, 2 mM MgCl2, 0.1 mM EDTA, 0.5 mM EGTA, 0.1 % NP-40, 150 mM KCl, 1x complete EDTA-free protease inhibitor (Roche) and 1x PhosStop (Roche) and 15 % glycerol). Cells were resuspended in binding buffer (100 OD_600_/ml). Bead beating was used to lyse the cell suspension until ∼80% cell lysis was obtained. Lysates were centrifuged at 15k rpm for 20 min, and the supernatant was collected. The extracted cell lysate was incubated with anti-FLAG M2 antibodies (Sigma) conjugated to Dynabeads™ M-280 sheep anti-mouse IgG (ThermoFisher Scientific) for 2 h at 4°C, under constant rotation. Unbound proteins were collected as flow-through and proteins bound to antibody-conjugated beads were washed five times with BB150 w/o glycerol, invert mixing was followed during each wash. Bound proteins were eluted in BB150 w/o glycerol + 200 μg/ml of 3x FLAG peptide (Sigma). Two elutes of ∼½ volume each of initial bead volume was taken and pooled.

1 μg of Anti-FLAG M2 antibody was conjugation to 10 μl of Dynabeads™ M-280 sheep anti-mouse IgG (ThermoFisher Scientific) in 1x phosphate-buffered saline (PBS), pH 7.4, and incubation for 1 h at room temperature (RT). Washed twice with 1x PBS and resuspended in PBS. This anti-FLAG conjugated beads were used for the lysate prepared from 100 OD_600_ culture.

Affinity purification samples that were processed subsequently for mass spectrometry was started from a 2.25 L culture, yielding ∼4500 OD_600_ cells. 300 mM KCl, where mentioned in experiments was used throughout the affinity purification experiment as part of the binding buffer yielding BB300.

For GFP affinity purification, GFP-Trap agarose beads (ChromoTek) were used. Bound proteins were eluted by boiling the beads for 10 min in 1x sample loading buffer (50 mM Tris-HCl pH 6.8, 2% SDS, 0.05% bromophenol blue, 10% glycerol, 5% 2-Metcaptoethanol) and the supernatant was collected. Other steps of the affinity purification protocol were kept the same as mentioned above.

For native-ChiP, lysate preparation, affinity purification, and isolation of the bound proteins were as mentioned above. DNA from the elute and input sample was extracted using MagExtractor clean-up kit (TOYOBO). PCR for the identical dilution of input and IP was set-up using centromere 14 primers (5’-GGTGATGCTACCTCGGT-3’ and 5’-CCCGACGACTGTATCAGTTA-3’) and non-centromere control primers (5’-GATCAAGTATAGGCGAAGG-3’ and 5’-CATCTCTTATTCCCACTTCTACTC-3’) located on the gene body of CNAG_00063, located ∼825 kb away from the centromere on chromosome 1 (Fig. 2a).

### Immunoblot analysis

For whole-cell lysates, 3 OD_600_ cells were harvested and resuspended in 15% TCA overnight. 500 μl 0.5 mm glass beads were added, and samples were vortexed for a total time of 15min, with intermittent cooling on ice. Centrifuged at 13k rpm for 10 min and the obtained pellet was washed twice with 100% acetone, air-dried and resuspended in 1x sample loading buffer and boiled for 10min. Samples were separated on an SDS-PAGE and transferred to Immobilon-P (Merck).

For Supplementary Fig. 1c and 5b and e, primary antibody and secondary antibody dilutions were made in skim milk. Proteins bound by antibodies were detected with Clarity western ECL (BioRad) and visualized with Versadoc (BioRad). For Figure 5c and Supplementary Fig. 1a and 4k, primary and secondary antibody dilution were prepared in Signal Enhancer Hikari (Nacalai tesque). ChemiDoc Touch (Bio-Rad) was used to visualize proteins reacting with antibody in the presence of the substrate ECL Prime (GE Healthcare). ImageJ^74, 75^ and Image lab (BioRad) was used to visualize and process images. Antibodies used are tabulated in Supplementary Table 8.

### Mass spectrometry

Affinity purified samples were separated on an SDS-PAGE followed by silver staining. Isolated samples from the stained gel were Trypsin digested. Samples were subject to nano LC-MS-MS as described previously^76^. Using the MASCOT ver2.6.2 search engine in Proteome Discoverer 2.1.1.21 and 2.2.0.388 (ThermoFisher Scientific) the obtained spectra peaks were assigned using the UniProt proteome database for *C. neoformans* H99α database (ID: UP000010091 20171201downloaded (7340 sequences)). Fragment tolerance 0.80 Da (Monoisotropic), parent tolerance 10 PPM (Monoisotropic), fixed modification of +57 on C (Carbamidomethyl), variable modification of +16 on M (oxidation) and +42 on Peptide amino-terminus (Acetyl) and allowing for a maximum of 2 missed cleavages for CENP-C^Mif2^, Dsn1 and Spc25 and 3 missed cleavages for bridgin samples. The obtained results were visualized using Scaffold 4.8.9 (Proteome Software). A minimum threshold for peptide (95%), and protein (99%) in addition to the identification of a minimum of two unique peptides were considered as hits after normalization with untagged control spectra. Identified protein hits from CENP-C^Mif2^, Dsn1, Spc25, and their untagged controls can be found in Supplementary Table 2. To relatively quantitate protein abundance obtained within each of the experiments Bgi1^FL^ 150 mM, Bgi1^FL^ 300 mM, and Bgi1^BDΔ^ 150 mM, emPAI^77^ (exponentially modified protein abundance index) values were determined using Scaffold 4.8.9 (Proteome Software). Higher the emPAI score, more abundant the protein is in the mixture. Supplementary Table 4 summarizes the identified interacting protein hits from Bgi1^FL^, Bgi1^BDΔ^, and their untagged control IPs.

### Cross-linked chromatin immunoprecipitation and quantitative real-time PCR

ChIP assays were performed with some modification of previously described protocols^78, 79^. In brief, 100 ml of Bgi1-GFP strain was grown until ∼1 OD_600_. Cross-linking was performed for 20min using formaldehyde to a final concentration of 1% and incubated at RT with intermittent mixing. The reaction was quenched by the addition of 2.5M glycine and further incubated for 5 min. Fixed cells were harvested by centrifugation and resuspended in 9.5 ml of deionized water, followed by the addition of ml of 2-Mercaptoethanol and incubated at 30°C for 60 min at 180 rpm. Cells were pelleted and resuspended in 10 ml spheroplasting buffer (1 M sorbitol, 0.1 M sodium citrate, and 0.01 M EDTA) containing 40 mg of lysing enzyme from *Trichoderma harzianum* (Sigma). Spheroplasts were washed once with 15 ml each of the following buffers, 1) 1x PBS 2) Buffer I (0.25% Triton X-100, 10 mM EDTA, 0.5 mM EGTA, 10 mM Na-HEPES pH 6.5) and 3) Buffer II (200 mM NaCl, 1 mM EDTA, 0.5 mM EGTA, 10 mM Na-HEPES pH 6.5). Following which the spheroplasts were resuspended in 1 ml of extraction buffer (50 mM HEPES pH 7.4, 1% Triton X-100, 140 mM NaCl, 0.1% Na-Deoxycholate, 1 mM EDTA) and sonicated to shear chromatin using a Bioruptor (Diagenode) for 30 cycles of 30 s on and 30 s off bursts at high-intensity setting. Sheared chromatin was isolated in the supernatant fraction after centrifugation for 15 min at 13k rpm. Average chromatin fragment sizes ranged from 200-500 bp. 100 μl, 1/10^th^ the volume, of the chromatin fraction, was kept for input DNA preparation, the remaining chromatin volume was divided into two halves of 450 μl each for (+) antibody and (-) antibody. For (+) antibody 20 μl of GFP-Trap agarose beads (ChromoTek) and 20 μl of blocked agarose beads (ChromoTek) was added to (-). The tubes were incubated for 8 h to overnight on a rotator at 4°C. Following which the supernatant was isolated as flow-through, and the beads were washed twice with low salt buffer, twice with high salt buffer, once with LiCl buffer and twice with TE. Bound chromatin was eluted in two 250 μl elution using elution buffer. All three fractions (SM, (+Ab) and (-Ab)) were de-crosslinked (mixed with 20 μl of 5 M NaCl and incubated at 65°C for 8 h to overnight), Proteinase K treated (10 μl of 0.5M EDTA, 20 μl of 1 M Tris-HCl pH 6.8, 40 mg Proteinase K was added to the solution and incubated for up to 2 h at 45°C) and DNA was isolated using phenol: chloroform extraction followed by ethanol precipitation. Isolated DNA was air-dried and dissolved in 25 μl of de-ionized water containing 25 μg/ml of RNase (Sigma).

All three samples (SM, (+) and (-) antibody) were subject to Real-time quantitative PCR. The reaction mixture was set up using the iTaq™ universal SYBR green Supermix (BioRad) with 1 μl of the undiluted (+Ab), (-Ab) DNA samples and SM (diluted 1:50). CN1 (CEN 14)-5’-CCATCCAGTTCTTGCTTGAG-3’ 5’-GCAAGGAATGTGTTGTCTGG-3’ and CN3 (CEN 2)-5’-CAGACCCTTCCTTCAGCCG-3’ 5’-TGGCAAGGAGTCGTCAGCG-3’ was used to estimate centromeric enrichment levels and non-centromeric primer set NC3 5’-GATCAAGTATAGGCGAAGG-3’ 5’-ATCTCTTATTCCCACTTCTACTC-3’ located ∼825 kb away from the centromere on chromosome 1 was used to normalize and obtained fold enrichment. Values were plot using GraphPad Prism.

### Fluorescence microscopy and analysis

Overnight cultures grown in YPD were sub-cultured into fresh YPD at 0.1 OD_600_ and grown until 0.4-OD_600_. Cells were isolated, washed twice in 1x PBS and mounted on slides. Images for Supplementary Fig. 1g (CNAG_01340) were acquired using the Airyscan mode in the Zeiss LSM 880 confocal system equipped with an Airyscan module, 63x Plan Apochromat 1.4 NA. Z-stacks were obtained at an interval of 166 nm, 488/516 and 561/595 nm excitation/emission wavelengths were used GFP and mCherry respectively. Airyscan images were processed using Zen (Zeiss) and visualized in ImageJ^74, 75^. Images for Figure 1f and Supplementary Fig. 1g and d were acquired in the Zeiss LSM 880 confocal system equipped with GaAsp photodetectors. Z-stacks were obtained at an interval of 300 nm, 488 nm and 561 nm excitation was used for GFP and mCherry respectively and emission between 490-553 nm and 571-651 nm was captured. Images are represented as maximum-intensity projections.

Live-cell microscopy, images for kinetochore quantitation and microscopy-based assays were acquired using the Zeiss Axio Observer 7, equipped with Definite Focus.2, Colibri 7 (LED light source), TempController 2000-2 (PECON), 100x Plan Apochromat 1.4 NA objective, pco.edge 4.2 sCMOS and Andor iXon Ultra 897 electron-multiplying CCD (charge-coupled device). Zen 2.3 (blue edition) was used for image acquisition and controlling all hardware components. Filter set 92 HE with excitation 455-483 nm and 583-600 nm for GFP and mCherry respectively, and corresponding emission was captured at 501-547 nm and 617-758 nm. To limit the time taken for an image, a complete Z-stack was obtained for each channel before switching.

For live-cell microscopy, an overnight culture was grown in YPD was sub-cultured into fresh YPD at ∼0.1 OD_600_ and grown for 2-3 generations until 0.4-0.8 OD_600_. Cells were harvested, washed in 1x PBS and resuspended in synthetic complete media with 3% dextrose. Cells were mounted onto an agarose pad (3% dextrose, 3% agarose in synthetic complete media) and sealed with petroleum jelly. Images were captured at time intervals of 0.5,1, 2 or 4 min, as appropriate, with an EM gain of 300 and Z interval of 300 nm. Z-stack projection of images are represented.

To study kinetochore interdependency, conditional strains were grown overnight in YPG, sub-cultured at 0.2 OD_600_ and grown until 0.8-1 OD_600_. Cells were washed and resuspended in 1x PBS. Following which cells were inoculated into YPD (repressive) and YPG (permissive) at 0.1 OD_600_. Images were acquired after 6, 12, 15, 18, 9 and 18 h for CENP-C^Mif2^, Mis12^Mtw1^, Nuf2, Knl1^Spc105^, Dad1, and Dad2 respectively. Z-stack was obtained at an interval of 300 nm. Single Z slice representing the maximum intensity of the tagged kinetochore proteins was represented. Quantitation of kinetochore signal was performed from large budded cells (budding index ∼0.55-0.90).

To estimate the population of large-bud and cells with segregation defects, cells were grown until early-log phase 0.8-1 OD_600_ after sub-culture from an overnight culture. Imaged using the above mentioned sCMOS camera with a Z-interval of 300 nm. Cells with a budding index of >0.55 were considered as large bud cells in mitosis. Chromatin marked with a tagged H4 construct was used to observe missegregation events.

Images for the over-expression assay of bridgin stains are representative maximum intensity projection images.

For live-cell quantitation of kinetochore signal, signal intensity was measured after the projection of Z-stacks. Kinetochore signal measurement in interdependency assays and *bgi1Δ* background were measured from the in-focus Z plane exhibiting the most intense signal. Background signal measured from a region neighboring the kinetochore measured signal in the same plane of the equal area was subtracted from the measured kinetochore intensity and normalized to the appropriate control and plot using GraphPad Prism 5.00 (GraphPad software). All acquired images were processed in ImageJ^74, 75^.

For images wherein, brightness and contrast are modified the settings were applied uniformly across the entire image.

### Budding index calculation

Budding index of a cell is defined as the ratio obtained by:

> *Diameter of daughter cell/Diameter of mother cell*

The diameter of the daughter and mother cell was measured along the mother-daughter axis using the line tool in ImageJ^75^

### Generation of recombinant proteins

GST, GST-Bgi1^FD^ (residues 1-130) and GST-Bgi1^BD^ (residues 1000-1295) was expressed from pGEX-6P-1 (GE Healthcare) in Rosetta2 (DE3) (Merck). GST and GST-Bgi1^BD^ were induced for expression using 1 mM IPTG for 3 h at 37°C. GST-FD was induced for expression overnight at 16°C using 0.2 mM IPTG. Cells were harvested and lysed in lysis buffer (20 mM HEPES pH 7.5, 300 mM NaCl, 1 mM EDTA, 0.5 mM TECP, 1x complete EDTA-free protease inhibitor (Roche)) and 1x PBS with 1x complete EDTA-free protease inhibitor (Roche) for GST and GST-Bgi1^FD^. GST fusion proteins were affinity purified using Glutathione sepharose 4b beads (GE Healthcare) and eluted using 20 mM glutathione. GST-Bgi1^FD^ and GST-Bgi1^BD^ were further purified using anion exchange chromatography. The column was equilibrated using 20 mM Tris-HCl pH 7.5, 1 mM DTT. Elution gradient of 5-75% NaCl was achieved using 20 mM Tris-HCl pH 7.5, 1 M NaCl, 1 mM DTT. Relevant fractions were pooled, concentrated in Amicon-Ultra (Merck), frozen in liquid nitrogen and stored at −80°C.

His-Bgi1^FL^ was expressed in SF9 cells. Cells were resuspended and lysed in binding buffer (20 mM Tris-HCl pH 8.0, 500 mM NaCl, 5 mM Imidazole). His-Bgi1^FL^ was affinity purified with Ni-NTA agarose (GE Healthcare), eluted with 20 mM Tris-HCl pH 8, 500 mM NaCl, 500 mM Imidazole. Purified protein was dialyzed against buffer containing Tris-HCl pH 7.5, 1 mM DTT and 100 mM NaCl. Samples were concentrated using Amicon-Ultra (Merck), frozen in liquid nitrogen and stored at −80°C. Absence of contaminating DNA was confirmed in all recombinant protein samples.

### Viability assay

An overnight culture was inoculated into fresh YPD medium at 0.1 OD_600_ and grown to ∼0.8 OD_600_. The cell number was measured, followed by dilution of the cell suspension. 100-500 cells were subsequently plated on YPD solidified using 2% agar and grown for 2 days at 30°C. The number of colonies formed was measured and plot as normalized values to the WT strain.

### Serial dilution growth analysis

Cells were grown overnight, inoculated into fresh YPD at 0.2 OD_600_ and grown until 0.8-1 OD_600_. Following which cells were isolated and made up to 2 OD/ml in 1x PBS. Further dilutions were made as indicated in 1x PBS. 2 μl of the cell suspension was transferred onto appropriate agar plates as mentioned and incubated for 2 days for 30°C and 30°C + TBZ control and 2 μg/ml TBZ, 3 days for 30°C + 4 μg/ml TBZ and 37°C and 7 days for 14°C.

### Electrophoretic mobility shift assays

Purified recombinant proteins of mentioned molar ratio were incubated with 601 DNA (2.5 pMoles) or 1 pMole of reconstituted nucleosomes in binding buffer (20 mM Tris pH 7.5, 100 mM NaCl, 5% glycerol and 1 mM DTT). Incubated for 1 h at 4°C and separated on a PAGE gel, stained with GelRed and visualized using a gel documentation system. Further, the gels were stained with Coomassie to visualize the protein complexes and imaged using a scanner.

### Estimation of DNA methylation

Genomic DNA was isolated from overnight cultures of WT and *bgi1Δ*, using a modified glass bead protocol^73^. In brief, cells were suspended in a microfuge tube containing 500 μl of lysis buffer (50 mM Tris-HCl pH 7.5, 20 mM EDTA and 1% SDS) and 250 μl of glass beads. Cells were disrupted by vortexing for 5 min and centrifuged for 1 min at 13k rpm. To the supernatant 275 μl of 7 M ammonium acetate was added and incubated at 65°C for 5 min and rapidly chilled on ice for 5 min. 500μl of chloroform was added, mixed and centrifuged at 13k rpm for 3 min. The supernatant, containing DNA was precipitated with isopropanol, washed with 70% ethanol, dried and resuspended in 50 μl deionized water.

The isolated genomic DNA was digested separately with CpG methylation-sensitive (HhaI) or insensitive (HindIII) restriction enzymes overnight with a no enzyme (uncut) control reaction. The digested DNA was diluted 1:50 and used for PCR amplification. Primer sets for PCR amplification of the centromeric region (5’-AGTCTCGTGTGGCTATGATT-3’ and 5’-GGATCTGCTTGACAGTGTCA-3’) and non-centromeric regions (5’-CCAACCGAAGCCCAAGACAA-3’ and 5’-TTGAAGGATGATCCGGCCGA-3’) were used. Obtained PCR products were subsequently separated by agarose gel electrophoresis using a 1% agarose gel and visualized by EtBr staining.

### Statistics and reproducibility

*P* values were assessed by unpaired, two-tailed *t-*test using GraphPad Prism 5.00 (GraphPad software). Error bars represent standard deviation (s.d.) or standard error of the mean (s.e.m.) as mentioned for each experiment. *N* for each experiment is mentioned in the figure legends.

## ACKNOWLEDGMENTS

The authors thank the members of the Sanyal and Fukagawa lab for inputs and discussions. We thank Daniel Gerlich (Institute of Molecular Biotechnology, Vienna, Austria) for generously providing us the Ki-67 plasmids. The authors are grateful to Masatoshi Hara, Mariko Ariyoshi, Reito Watanabe, and Fumiaki Makino from the Fukagawa lab for their technical assistance. We also thank Akira Shinohara and his lab members for allowing us to use their lab facilities. A joint grant from the Department of Science and Technology, India (DST) and Japanese Society for the Propagation of Science, Japan (JSPS) aided in the travel of S.S. between the Sanyal and Fukagawa labs. Research in the Sanyal lab was funded by intramural funding from JNCASR. S.S. thanks Council for Scientific and Industrial Research (CSIR) for his fellowship.

## AUTHOR CONTRIBUTIONS

S.S., K.S., T.H., and T.F. designed the experiments. S.S. performed the experiments. T.H. assisted with affinity purification for mass spectrometry, and R.N. performed the mass spectrometry. S.S., K.S., and T.F. wrote the manuscript with discussion and inputs from all the authors.

