## Supplementary table 6 for "Identification of bridgin, an unconventional linker, connects the outer kinetochore to centromeric chromatin"

**Supplementary table 6: Strains and plasmids**

**Strain used in this study:**

| Strain name | Genotype | Reference |
|---|---|---|
| H99α | *MATα (Wild type)* | 1 |
| KN99a | *MATa (Wild type)* | 2 |
| SHR896 | *MATα CENP-C$^{MIF2}$::CENP-C$^{MIF2}$-3xFLAG-HygB* | This study |
| SHR824 | *MATα DSN1::DSN1-3xFLAG-HygB, NUF2::NUF2-GFP-NAT, Mis12$^{MTW1}$:: Mis12$^{MTW1}$-mCherry-NEO* | This study |
| SHR861 | *MATα SPC25::SPC25-3xFLAG-HygB* | This study |
| SHR823 | *MATα SPC25::SPC25-3xFLAG-HygB, NUF2::NUF2-GFP-NAT, Mis12$^{MTW1}$:: Mis12$^{MTW1}$-mCherry-NEO* | This study |
| SHR845 | *MATa SOS7::GAL7p-SOS7-HYGB, K*99::mCherry-CENP-A-NEO (pLKB74) | This study |
| SHR876 | *MATa BGI1::BGI1-V5-GFP-NAT, CENP-C$^{MIF2}$:: CENP-C$^{MIF2}$-mCherry-NEO* | This study |
| SHR897 | *MATa CENP-C$^{MIF2}$:: CENP-C$^{MIF2}$-mCherry-NEO, BKT2::BKT2-V5-GFP-NAT* | This study |
| SHR842 | *MATa CENP-C$^{MIF2}$:: CENP-C$^{MIF2}$-mCherry-NEO, YTA7::YTA7-V5-GFP-NAT* | This study |
| SHR905 | *MATa KN*99::mCherry- *CENP-A$^{CSE4}$*-NEO (pLKB74), *MCM6::MCM6-V5-GFP-NAT* | This study |
| SHR870 | *MATα BGI1::BGI1-V5-GFP-NAT* | This study |
| SHR516 | *MATa Mis12$^{MTW1}$:: Mis12$^{MTW1}$-mCherry-NEO, NUF2::NUF2-GFP-NAT* | 3 |
| CNVY120 | *MATa KN99::GFP-DAD1-NAT (pVY2), KN99:: CENP-A$^{CSE4}$-mCherry-NEO* | 3 |
| CNV119 | *MATα H99::GFP-DAD1-NAT (pVY2), DAD2::DAD2-mCherry-NEO* | 3 |
| SHR772 | *MATa Mis12$^{MTW1}$:: Mis12$^{MTW1}$-mCherry-NEO, KNL1$^{SPC105}$:: KNL1$^{SPC105}$-GFP-NAT* | This study |
| SHR869 | *MATa Mis12$^{MTW1}$:: Mis12$^{MTW1}$-mCherry-NEO, BGI1::BGI1-V5-GFP-NAT* | This study |
| SHR906 | *MATα DAD2::GAL7p-mCherry-DAD2-HygB, BGI1::BGI1-V5-GFP-NAT* | This study |
| SHR907 | *MATa Mis12$^{MTW1}$::GAL7p-mCherry- Mis12$^{MTW1}$-HygB, BGI1::BGI1-V5-GFP-NAT* | This study |
| SHR908 | *MATα BGI1::BGI1-V5-GFP-NAT, sos7Δ::NEO* | This study |
| SHR909 | *MATa Dad2::DAD2-mCherry-NEO, BGI1::BGI1-V5-GFP-NAT* | This study |
| SHR720 | *MATa NDC80::NDC80-mCherry-NEO, CENP-C$^{MIF2}$::GAL7p-GFP- CENP-C$^{MIF2}$-HygB* | This study |
| SHR724 | *MATa NDC80::NDC80-mCherry-NEO, Mis12$^{MTW1}$::GAL7p-GFP-Mis12$^{MTW1}$-HygB* | This study |

| | | |
|---|---|---|
| SHR732 | *MATa Mis12*$^{MTW1}$*:: Mis12*$^{MTW1}$*-mCherry-NEO, NUF2::GAL7p-GFP-NUF2-HygB* | This study |
| SHR910 | *MATα KNL1*$^{SPC105}$*:: KNL1*$^{SPC105}$*-GFP-NAT, NUF2::GAL7p-mCherry-NUF2-HygB* | This study |
| SHR768 | *MATα KNL1*$^{SPC105}$*:: KNL1*$^{SPC105}$*-GFP-NAT, Mis12*$^{MTW1}$*::GAL7p-mCherry-Mis12*$^{MTW1}$*-HygB* | This study |
| SHR911 | *MATa DAD2::DAD2-mCherry-NEO, NUF2::GAL7p-GFP-NUF2-HygB* | This study |
| SHR767 | *MATα KNL1*$^{SPC105}$*:: KNL1*$^{SPC105}$*-GFP-NAT, Dad2::GAL7p-mCherry-DAD2-HygB* | This study |
| SHR713 | *MATa NDC80::NDC80-mCherry-NEO, DAD1::GAL7pDAD1* | This study |
| SHR867 | *MATα bgi1Δ::NEO* | This study |
| SHR838 | *MATa bgi1Δ::HygB* | This study |
| SHR830 | *MATα* H99::GFP-H4-NAT (pLKB35), *bgi1Δ::HygB* | This study |
| SHR832 | *MAT a H4::H4-mCherry-NEO, bgi1Δ::HygB* | This study |
| SHR873 | *MAT a H4::H4-mCherry-NEO, KN99::GFP-PCNA-NAT (pSS60)* | This study |
| SHR912 | *MATa bgi1Δ::HygB, KN99::3xFLAG-GFP-BGI1-NAT* | This study |
| SHR879 | *MATa H4::H4-mCherry-NEO, bgi1Δ::HygB, KN99::3xFLAG-GFP-BGI1-NAT (pSS62)* | This study |
| SHR854 | *MAT a H4::H4-mCherry-NEO, KN99::GFP-PCNA-NAT (pSS60)* | This study |
| SHR734 | *MATα mad2Δ::NEO,* | This study |
| SHR866 | *MATα mad2Δ::NEO, bgi1Δ::HygB, H4::H4-GFP-NAT* | This study |
| SHR741 | *MATa H4::H4-mCherry-NEO, mad2Δ::NEO* | This study |
| SHR913 | *MAT a H4::H4-mCherry-NEO, bgi1Δ::HygB, bgi1Δ::3x-FLAG-GFP-BGI1 FDΔ-NAT (pSS63)* | This study |
| SHR880 | *MAT a H4::H4-mCherry-NEO, bgi1Δ::HygB, bgi1Δ::3x-FLAG-GFP-BGI1 BDΔ-NAT (pSS64)* | This study |
| SHR915 | *MAT a H4::H4-mCherry-NEO, bgi1Δ::HygB, bgi1Δ::3x-FLAG-GFP-BGI1 FD-NAT (pSS65)* | This study |
| SHR916 | *MAT a H4::H4-mCherry-NEO, bgi1Δ::HygB, bgi1Δ::3x-FLAG-GFP-BGI1 USD-NAT (pSS66)* | This study |
| SHR917 | *MAT a H4::H4-mCherry-NEO, bgi1Δ::HygB, bgi1Δ::3x-FLAG-GFP-BGI1 BD-NAT (pSS67)* | This study |
| SHR918 | *MAT a H4::H4-mCherry-NEO, bgi1Δ::HygB, bgi1Δ::3x-FLAG-GFP-NAT (pSS61)* | This study |
| CNVY121 | *MAT a H4::H4-mCherry-NEO* | 3 |
| SHR903 | *MATa Mis12*$^{MTW1}$*:: Mis12*$^{MTW1}$*-mCherry-NEO, NUF2::NUF2-GFP-NAT, bgi1Δ::HygB* | This study |
| SHR904 | *MATa Mis12*$^{MTW1}$*:: Mis12*$^{MTW1}$*-mCherry-NEO, KNL1*$^{SPC105}$*:: KNL1*$^{SPC105}$*-GFP-NAT, bgi1Δ::HygB* | This study |

| | | |
|---|---|---|
| SHR902 | *MATa KN99::GFP-DAD1-NAT (pVY2), KN99::CENP-A$^{CSE4}$-mCherry-NEO, bgi1Δ::HygB* | This study |
| SHR919 | *MAT a bgi1Δ::HygB, bgi1Δ::3x-FLAG-FD-NAT (pSS77)* | This study |
| SHR858 | *MATa NDC80::NDC80-mCherry-NEO, BGI1::GAL7p-GFP-BGI1-HygB* | This study |
| SHR893 | *MATα SPC34::SPC34-3xFLAG-GFP-NAT* | This study |
| SHR895 | *MAT a H4::H4-mCherry-NEO, bgi1Δ::HygB, KN99::H3p-GFP-BGI1-NAT (pSS68)* | This study |
| SHR920 | *MAT a H4::H4-mCherry-NEO, bgi1Δ::HygB, KN99::H3p-GFP-BGI1-FDΔ-NAT (pSS69)* | This study |
| SHR921 | *MAT a H4::H4-mCherry-NEO, bgi1Δ::HygB, KN99::H3p-GFP-BGI1-BDΔ-NAT (pSS70)* | This study |
| SHR922 | *MAT a H4::H4-mCherry-NEO, bgi1Δ::HygB, KN99::H3p-GFP-BGI1-FD-NAT (pSS71)* | This study |
| SHR923 | *MAT a H4::H4-mCherry-NEO, bgi1Δ::HygB, KN99::H3p-GFP-BGI1-USD-NAT (pSS72)* | This study |
| SHR924 | *MAT a H4::H4-mCherry-NEO, bgi1Δ::HygB, KN99::H3p-GFP-BGI1-BD-NAT (pSS73)* | This study |
| SHR925 | *MAT a H4::H4-mCherry-NEO, KN99::H3p-GFP-BGI1-NAT (pSS74)* | This study |
| SHR926 | *MAT a H4::H4-mCherry-NEO, bgi1Δ::HygB, MAT a H4::H4-mCherry-NEO, bgi1Δ::HygB, bgi1Δ::3x-FLAG-GFP-(BDΔ+HsKi67BD)-NAT (pSS75)* | This study |

**Plasmid used in this study:**

| **Plasmid name** | **Description** | **Reference** |
|---|---|---|
| pCIN19 | *H3p-GFP-NAT* | Alspaugh lab |
| pSS59 | *BGI1-3xFLAG-NAT in pBlueScriptII KS(-)* | This study |
| pSS61 | *BGI1p-3xFLAG-GFP-NAT in pBlueScriptII KS(-)* | This study |
| pSS55 | *CENP-C$^{MIF2}$-3xFLAG-HygB in pBlueScriptII KS(-)* | This study |
| pSS56 | *DSN1-3xFLAG-HygB in pBlueScriptII KS(-)* | This study |
| pSS57 | *SPC25-3xFLAG-HygB in pBlueScriptII KS(-)* | This study |
| pSS78 | *BGI1-V5-GFP-NAT in pRS426* | This study |
| pSS85 | *GAL7p-GFP-SOS7-HygB in pBlueScriptII KS(-)* | This study |
| pSS79 | *BKT2-V5-GFP-NAT in pRS426* | This study |
| pSS81 | *BKT3-V5-GFP-NAT in pRS426* | This study |
| pSS80 | *YTA7-V5-GFP-NAT in pRS426* | This study |
| pSS82 | *MCM6-V5-GFP-NAT in pRS426* | This study |
| *pSS13* | *GAL7p-GFP- CENP-C$^{MIF2}$-NAT pBlueScriptII KS(-)* | This study |
| *pSS14* | *GAL7p-GFP- Mis12$^{MTW1}$-NAT pBlueScriptII KS(-)* | This study |

| | | |
|---|---|---|
| *pSS7* | *GAL7p-GFP-NUF2-NAT pBlueScriptII KS(-)* | This study |
| *pSS24* | *GAL7p-mCherry-NUF2-NAT pBlueScriptII KS(-)* | This study |
| *pSS19* | *GAL7p-GFP- Mis12$^{MTW1}$-NAT pBlueScriptII KS(-)* | This study |
| *pSS27* | *GAL7p-GFP-KNL1$^{SPC105}$-NAT pBlueScriptII KS(-)* | This study |
| *pSS21* | *GAL7p-mCherry-DAD2-NAT pBlueScriptII KS(-)* | This study |
| *pSS4* | *GAL7p-GFP-DAD1-NAT pBlueScriptII KS(-)* | This study |
| pSS84 | *BGI1 FL in pSS59 (BamHI/SpeI)* | This study |
| pSS62 | *BGI1 FL in pSS61 (BamHI/SpeI)* | This study |
| pSS63 | *BGI1 FDΔ in pSS61 (BamHI/SpeI)* | This study |
| pSS64 | *BGI1 BDΔ in pSS61 (BamHI/SpeI)* | This study |
| pSS65 | *BGI1 FD in pSS61 (BamHI/SpeI)* | This study |
| pSS66 | *BGI1 USD in pSS61 (BamHI/SpeI)* | This study |
| pSS67 | *BGI1 BD in pSS61 (BamHI/SpeI)* | This study |
| pSS76 | *BGI1 FL in pSS59 (BamHI/SpeI)* | This study |
| pSS58 | *SPC34-3xFLAG-HygB in pBlueScriptII KS(-)* | This study |
| pSS86 | *BGI1 FD in pGEX6P1* | This study |
| pSS87 | *BGI1 BD in pGEX6P1* | This study |
| pSS89 | *6xHis-BGI1 in pFASTBacHTA* | This study |
| pSS87 | *GAL7p-GFP-BGI1-HygB in pBlueScriptII KS(-)* | This study |
| pSS68 | *BGI1 FL (BamHI/SpeI) in pCIN19* | This study |
| pSS69 | *BGI1 FDΔ (BamHI/SpeI) in pCIN19* | This study |
| pSS70 | *BGI1 BDΔ (BamHI/SpeI) in pCIN19* | This study |
| pSS71 | *BGI1 FD (BamHI/SpeI) in pCIN19* | This study |
| pSS72 | *BGI1 USD (BamHI/SpeI) in pCIN19* | This study |
| pSS73 | *BGI1 BD (BamHI/SpeI) in pCIN19* | This study |
| pSS74 | *BGI1 HsKi67 BD (BamHI/SpeI) in pCIN19* | This study |
| pSS75 | *BGI1 BDΔ+HsKi67BD in pSS61 (BamHI/SpeI)* | This study |
