## Supplementary table 7 for "Identification of bridgin, an unconventional linker, connects the outer kinetochore to centromeric chromatin"

**Supplementary table 7: Primer list**

**Primers used in the study:**

| **Name** | **Sequence (5’------3’)** | **Description** |
|---|---|---|
| SHR3 | GTGCGAGCTCGCTAGCTTCTCCAAGATGGGTGTCACG | Generation of GAL7p-GFP-DAD1 |
| SHR4 | GTGAGAATGCGGCCGCCTTGGAGTGCTAGTTTTCCTGC | |
| SHR5 | ATGTCTTTATCAAGACCATCGAATGCCTACGATGC | |
| SHR6 | GTGCGGTACCGAGCTCATGCCTATGAAGTCCAGC | |
| SHR72 | AGCTTGAGCTCCTTCGAGATATACAGCTCC | Generation of GAL7p-GFP/mCherry-DAD2 |
| SHR73 | TTTAAGCGGCCGCCACTCGAGAGTTACAGTG | |
| SHR74 | CATCAAGCTTGGTGGTATGTCCCGTCCATCAATAGAAATG | |
| SHR75 | AACTCTCGAGGTGAGATAGGGTTGAAGGAGC | |
| SHR64 | AGCTGAGCTCCAAATCCACAACATCTGAAATACG | Generation of GAL7p-GFP/mCherry-Mis12$^{MTW1}$ |
| SHR65 | AAATTTGCGGCCGCGAACGTAGAGACGATTATGAATGC | |
| SHR66 | GCTGTTAACGGTGGTATGGTCCCGAGGAAGCCAG | |
| SHR67 | AGGTCCTCGAGCATTGGCAAGCTAACTAAATTAATGGAACG | |
| SHR68 | AGCTGAGCTCCAAGTCTCTTGTCGACATCTCTCC | Generation of GAL7p-GFP/mCherry-CENP-C$^{MIF2}$ |
| SHR69 | TTATTAGCGGCCGCGTTGAAGATGTTCTGGAGAAGTGC | |
| SHR70 | AACCCAAGCTTATGTCCCACATAACACCCTCAAGA | |
| SHR71 | TCGTCTCGAGCTTTCCATCTGCTTGCTTCTTTGG | |
| SHR50 | GACTGAGCTCCTTGCACTCTTACAGAAGCCTCC | Generation of GAL7p-GFP/mCherry-NUF2 |
| SHR51 | TCACATGCGGCCGCGATTGCTGAATGCAAATGCAG | |
| SHR52 | GACTAAGCTTATGTCGCAGCAGAATCGCAG | |
| SHR53 | GACTGGTACCGATTTCAAGCTGTGTGACGATACG | |
| SHR112 | AATGCGAGCTCTCTGTACCAGATAGTCACCAC | Generation of GAL7p-GFP-KNL1$^{SPC105}$ |
| SHR113 | ATATATATGCGGCCGC AATATGCTCGGTTAACTGCTG | |
| SHR114 | TAGTCAAGCTTATGTCTTTAGCAGCTCGCTC | |
| SHR115 | TCTAGGTACCGTTTCGAGTTGCTGTAGCTG | |
| SHR179 | CTACTCTTACAGGCAAGTTGGAG | KNL1$^{SPC105}$-GFP tagging |
| SHR180 | CTCGCCCTTGCTCACCATACTGGAGTACCTTGCACCGA | |
| SHR181 | TCGGTGCAAGGTACTCCAGTATGGTGAGCAAGGGCGAG | |
| SHR182 | CCTTGTAACCATCCATACAACCTAGGATGTGAGCTGGAGAGC | |
| SHR183 | GCTCTCCAGCTCACATCCTAGGTTGTATGGATGGTTACAAGG | |
| SHR184 | CTCTGGTGATACACTCAAGGAC | |
| SHR77 | TCGTGAGCTCGTCTCAACAATTTGGTTACTGATCAAGG | Generation of mad2Δ cassette |
| SHR78 | AGGACACTAGTTTCGTGGGGTAGAAACTGGAAG | |
| SHR81 | TTTAAAGCGGCCGCGTAATATTATCTAGTTCAACGTTCACG | |
| SHR80 | ACCCTTAGATCTGTGAATTCCTTTTATCCATTTTCC | |

| | | |
|---|---|---|
| SHR389 | GTCAGAGCTCCTCACAACATAAGACATCG | Generation of GAL7p-GFP-SOS7 |
| SHR390 | ATATATGCGGCCGCAGATCCAATATTACTACTATACGG | |
| SHR391 | GTCAAAGCTTGCTGGTGCAGGAATGGAACCCTCTATGACG | |
| SHR392 | GTCACTCGAGGTTTGAGCTTCAACCAG | |
| SHR374 | GATGTTGAGAGAAGTGATGGAGG | Tagging of DSN1 with 3xFLAG |
| SHR450 | CTACTTGTCATCGTCATCCTTGTAGTCGATGTCATGATCTTTATAATCACCGTCATGGTCTTTGTAGTCTTCCCTCTCCGGCCTA | |
| SHR452 | CGACTACAAGGATGACGATGACAAGTAGCTAGTAACGGCCGCCA | |
| SHR377 | CAATTGTAACCATCGTCATTAACACCAGTGTGATGGATATCTGCAGA | |
| SHR378 | TCTGCAGATATCCATCACACTGGTGTTAATGACGATGGTTACAATTG | |
| SHR379 | GATGGCATTTCGCTAACCAC | |
| SHR367 | GTATGTGTCGACGTATGACCT | |
| SHR451 | CTACTTGTCATCGTCATCCTTGTAGTCGATGTCATGATCTTTATAATCACCGTCATGGTCTTTGTAGTCTTTACCCAAAGCCAATTG | Tagging of SPC25 with 3xFLAG |
| SHR370 | GCACTCAAAAATGTTACAAATACAGTCCAGTGTGATGGATATCTGCAGA | |
| SHR371 | TCTGCAGATATCCATCACACTGGACTGTATTTGTAACATTTTTGAGTGC | |
| SHR372 | CATCGTCATGCCAATCGTG | |
| SHR493 | GAAGAATGGTAGAGCAAGG | Tagging of CENP-C$^{MIF2}$ with 3xFLAG |
| SHR511 | CTTGTCATCGTCATCCTTGTAGTCGATGTCATGATCTTTATAATCACCGTCATGGTCTTTGTAGTCTCCAGCACCTCTCCTACTCTTCCCCTTAC | |
| SHR494 | ACCCATTCATACCTTCTTTCTCAGTGTGATGGATATCTGCAGA | |
| SHR495 | TCTGCAGATATCCATCACACTGAGAAAGAAGGTATGAATGGGT | |
| SHR496 | CACCAGATAGAAAGAGTCTAGG | |
| SHR513 | CGACGGTATCGATAAGCTTGATATCGAGATGTACGAGGAAGAAGAGG | Tagging of BGI1 with V5-GFP |
| SHR514 | GAGACCAAGGAGAGGGTTGGGGATAGGCTTACCAGCACCCTTCCTACTCCTGGTTGTCCT | |
| SHR515 | GCACCTATCTTACAACATCCACTATCAGGATGTGAGCTGGAGAGC | |
| SHR516 | GCTCTCCAGCTCACATCCTGATAGTGGATGTTGTAAGATAGGTGC | |
| SHR517 | CGGCCGCTCTAGAACTAGTCAGAGGAAGGAACCTTGGATG | |

| | | |
|---|---|---|
| SHR518 | CGACGGTATCGATAAGCTTGATATCCAATGGAGCTCTCCAGATGTC | Tagging of YTA7 with V5-GFP |
| SHR519 | GAGACCAAGGAGAGGGTTGGGGATAGGCTTACCAGCACCATCGTTTTTCCAACTATTAACCTCTTTG | |
| SHR520 | AAACGCCATGCTAACAACAAAATGAGGATGTGAGCTGGAGAGC | |
| SHR521 | GCTCTCCAGCTCACATCCTCATTTTGTTGTTAGCATGGCGTTT | |
| SHR522 | CGGCCGCTCTAGAACTAGTCTCCATCTTCGTTCATTCACGC | |
| SHR523 | CGACGGTATCGATAAGCTTGATATCCAGTCAGGGAAGATTTGACGTG | Tagging of BKT2 with V5-GFP |
| SHR524 | GAGACCAAGGAGAGGGTTGGGGATAGGCTTACCAGCACCCTCGTCACCACCGAACAC | |
| SHR525 | GCATTAGTGTGGCTTCTTGATTCAGGATGTGAGCTGGAGAGC | |
| SHR526 | GCTCTCCAGCTCACATCCTGAATCAAGAAGCCACACTAATGC | |
| SHR527 | CGGCCGCTCTAGAACTAGTCATTCAAGGTAGCACATAAAGTTGAC | |
| SHR533 | CGACGGTATCGATAAGCTTGATATCACTGCTGAGAGGAGCTGTG | Tagging of BKT3 with V5-GFP |
| SHR534 | GAGACCAAGGAGAGGGTTGGGGATAGGCTTACCAGCACCGATCATTTCGTAACCTTCATCTTTTGC | |
| SHR535 | CGCTATACTACCTTAAGTTTAACCGTAGGATGTGAGCTGGAGAGC | |
| SHR536 | GCTCTCCAGCTCACATCCTACGGTTAAACTTAAGGTAGTATAGCG | |
| SHR537 | CGGCCGCTCTAGAACTAGTCCAACACACAAATTATCAAGGATTCC | |
| SHR453 | CGACGGTATCGATAAGCTTGATATCGATCTTCGTCAGCATCTAGCTC | Tagging of SPC34 with 3xFLAG-GFP |
| SHR454 | TATAATCACCGTCATGGTCTTTGTAGTCAGCTCCATCTGCAAATCTAACCCTACCC | |
| SHR455 | GGTATACAGTTAGATCAAGGAGGATACAGGATGTGAGCTGGAGAGC | |
| SHR456 | GCTCTCCAGCTCACATCCTGTATCCTCCTTGATCTAACTGTATACC | |
| SHR457 | CGGCCGCTCTAGAACTAGTCAAATAACATGACGTGACGGAC | |
| SHR538 | CGACGGTATCGATAAGCTTGATATCGCTCCAGAGGTATATTCGATACG | Tagging of MCM6 with V5-GFP |
| SHR539 | GAGACCAAGGAGAGGGTTGGGGATAGGCTTACCAGCACCTGCGGGAATAGAAGAAGATAAATCTG | |
| SHR540 | GGAACAGCGGGAAATGCAAGGATGTGAGCTGGAGAGC | |

| | | |
|---|---|---|
| SHR541 | GCTCTCCAGCTCACATCCTTGCATTTCCCGCTGTTCC | |
| SHR542 | CGGCCGCTCTAGAACTAGTCGAACCCTGCTCAAGTCG | |
| SHR560 | GGTAAGCCTATCCCCAACCCTCTCCTTGGTCTCGACAGCACCGGTGCTATGGTGAGCAAGGGCGAG | Common V5-GFP primer |
| SHR548 | GCTCAGAGGTCACATACAGG | Generation of sos7Δ cassette |
| SHR564 | CTGCAGATATCCATCACACTGGAGGTCAAAGATGGGTAAATAGC | |
| SHR565 | GCTATTTACCCATCTTTGACCTCCAGTGTGATGGATATCTGCAG | |
| SHR551 | GCTGTCCACTTTCGAAGGTCAGTGTGCTGGAATTCGC | |
| SHR552 | GCGAATTCCAGCACACTGACCTTCGAAAGTGGACAGC | |
| SHR553 | CATTATTGGAGATGTCTGAAGCG | |
| SHR582 | CGACGGTATCGATAAGCTTGATATCCCAGAAGGATAGAGTCCTCTG | Generation of GAL7p-GFP/mCherry-BGI1 |
| SHR583 | CTCACATCCTCGCAGCTTTTCGTTGCAAGTCAGC | |
| SHR584 | GCTGACTTGCAACGAAAAGCTGCGAGGATGTGAG | |
| SHR585 | CTCTCGTCAAACTCTTGCATGGCACCAGCGTACAGCTCGTCCATGCCG | |
| SHR586 | CGGCATGGACGAGCTGTACGCTGGTGCCATGCAAGAGTTTGACGAGAG | |
| SHR587 | CGGCCGCTCTAGAACTAGTGTTACTGTCAATTGAGGAAGC | |
| SHR600 | CTAGCTTGGCAATAGTGTAGCAG | Generation of *bgi1Δ* cassette |
| SHR601 | CTGCAGATATCCATCACACTGGGTTGCTGTTTGTATAGCGAGTC | |
| SHR602 | GACTCGCTATACAAACAGCAACCCAGTGTGATGGATATCTGCAG | |
| SHR603 | GCACCTATCTTACAACATCCACTATCCAGTGTGCTGGAATTCGC | |
| SHR604 | GCGAATTCCAGCACACTGGATAGTGGATGTTGTAAGATAGGTGC | |
| SHR605 | CAGAGGAAGGAACCTTGGATG | |
| SHR611 | AGTCGGATCCGCCGCTGGTGCCATGCAAGAGTTTGACGAGAG | Generation of domain deletion constructs of BGI1 expressed under native or H3 promoter |
| SHR612 | ATATATACTAGTCAAGTACTCGCCACTTATCACTC | |
| SHR613 | AGTCGGATCCGCCGCTGGTGCCTGGACCTATGGAAGATGCT | |
| SHR614 | ATATATACTAGTCTATCTTACAACATCCACTATCCATTATCATTAAGCATCTTCCATAGGTCCA | |
| SHR615 | ATATATACTAGTCTATCTTACAACATCCACTATCCATTATCATTATAGTTGAACCTGAAAAGCTTTTTC | |
| SHR616 | AGTCGGATCCGCCGCTGGTGCCGAGATCGAAGAGAAGGGTAAAG | |
| SHR715 | CCTCAGTGTGGCCTGGGGCACCAGCCTCATCTTGCTCCTGCAC | |

| | | |
|---|---|---|
| SHR716 | GTGCAGGAGCAAGATGAGGCTGGTGCCCCAGGCCACACTGAGG | Generation of fusion protein of BGI1-BDΔ and HsKi67-BD |
| SHR676 | ATATATACTAGTCTATCTTACAACATCCACTATCCATTATCATTACCAAATATCTTCACTGTCCCTATG | |
| SHR677 | AGTCGGATCCCCAGGCCACACTGAGG | Over-expression of HsKi67BD |
| SHR729 | CTGAGGATCCATGCAAGAGTTTGACGAGAG | Tagging BGI1-FD with GST for recombinant protein purification |
| SHR730 | CTGAGTCGACTTAATCTTCCATAGGTCCATAGTTG | |
| SHR731 | CTGAGGATCCATCGAAGAGAAGGGTAAAGG | Tagging BGI1-BD with GST for recombinant protein purification |
| SHR733 | ATATATGCGGCCGCTTACTTCCTACTCCTGGTTGTC | |
| VYP75 | AGTCTCGTGTGGCTATGATT | Centromeric DNA methylation |
| VYP76 | GGATCTGCTTGACAGTGTCA | |
| VYP79 | CCAACCGAAGCCCAAGACAA | Non-centromeric DNA methylation |
| VYP80 | TTGAAGGATGATCCGGCCGA | |
