## Supplementary table 8 for "Identification of bridgin, an unconventional linker, connects the outer kinetochore to centromeric chromatin"

**Supplementary table 8: List of antibodies used in the study**

**Primary antibodies**

| Antibody | Species | Source | Catalogue no | Assay and dilution |
|---|---|---|---|---|
| α-H3K9me2 | Mouse | Abcam | ab1220 | Immunoblot, 1:2000 |
| α-PSTAIR | Mouse | Abcam | 10345 | Immunoblot, 1:5000 |
| α-GFP | Mouse | Roche | 11814460001 | Immunoblot, 1:3000 |
| α-H4 | Mouse | Reference 1 | CMA400 | Immunoblot, 1:5000 |
| α-FLAG M2 | Mouse | Sigma | F3165 | Immunoblot, 1:5000 |
| α-pan H3 | Rat | Reference 2 | 140-1G1 | Immunoblot, 1:3000 |

**Secondary antibodies**

| Antibody | Species | Source | Catalogue no | Assay and dilution |
|---|---|---|---|---|
| α-Mouse | Goat | Bangalore genei | HO06 | Immunoblot, 1:10000 |
| HRP-conjugated α-mouse IgG | Rabbit | Jackson ImmunoResearch | 315-035-003 | Immunoblot, 1:15000 |
| HRP-conjugated α-Rat IgG | Goat | Jackson ImmunoResearch | 112-035-003 | Immunoblot, 1:15000 |


1. Hayashi-Takanaka, Y. *et al.* Distribution of histone H4 modifications as revealed by a panel of specific monoclonal antibodies. *Chromosome Res.* 23, 753-766 (2015).
2. Kimura, H., Hayashi-Takanaka, Y., Goto, Y., Takizawa, N. & Nozaki, N. The organization of histone H3 modifications as revealed by a panel of specific monoclonal antibodies. *Cell Struct Funct.* 33, 61-73 (2008).
