## Supplementary table 3 for "Identification of bridgin, an unconventional linker, connects the outer kinetochore to centromeric chromatin"

**List of bridgin repeats**

| Start | *p*-value | | sequence | |
|---|---|---|---|---|
| 2 | 1.21E-04 | E | EEDEVQ | E |
| 2 | 1.21E-04 | E | EEDEVQ | E |
| 2 | 1.21E-04 | E | EEDEVQ | E |
| 4 | 2.93E-04 | DEY | EEYEVQ | |
| 2 | 1.15E-03 | E | EEEEVQ | E |
| 2 | 1.15E-03 | E | EEEEVQ | E |
| 2 | 1.95E-03 | E | EEKEVD | L |
| 2 | 2.14E-03 | E | EEKESD | D |
| 2 | 8.67E-03 | E | EKEERN | P |
| 3 | 1.20E-02 | TE | EEEEYL | |
| 1 | 4.36E-02 | | EEEEEQ | KE |
| 1 | 4.93E-02 | | EEEGVE | EL |
| 2 | 6.16E-02 | E | VEEERK | P |
