## Supplementary information for "Identification of bridgin, an unconventional linker, connects the outer kinetochore to centromeric chromatin"

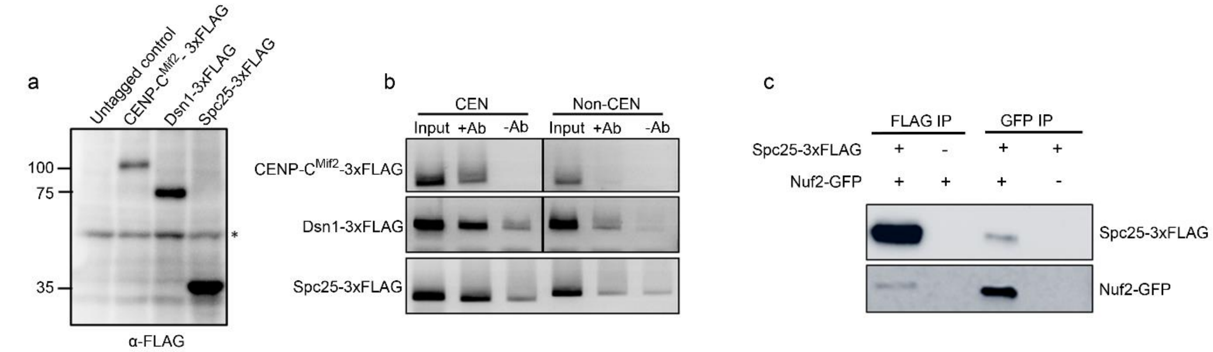


d Proteins of unknown function

| Orf | Given name | CENP-C$^{Mif2}$-3xFLAG % coverage | CENP-C$^{Mif2}$-3xFLAG peptides | Dsn1-3xFLAG % coverage | Dsn1-3xFLAG Peptides | SPC25-FLAG % coverage | SPC25-FLAG Peptides | Domains present | Domain function |
|---|---|---|---|---|---|---|---|---|---|
| CNAG_01903 | Bkt1 | 46 | 120 | 46 | 120 | 47 | 212 | FHA domain | Phosphopeptide recognition |
| CNAG_03959 | Bkt2 | 65 | 17 | 65 | 17 | 73 | 27 | NA | NA |
| CNAG_02701 | Bkt3 | 17 | 10 | 17 | 10 | 15 | 6 | BAG domain | Modulators of chaperone activity |

e Predicted chromatin binding proteins with no known function at the kinetochore

| Orf | Given name | CENP-C$^{Mif2}$-3xFLAG % Coverage | CENP-C$^{Mif2}$-3xFLAG peptides | Dsn1-3xFLAG % coverage | Dsn1-3xFLAG Peptides | SPC25-FLAG % coverage | SPC25-FLAG Peptides | Domains present |
|---|---|---|---|---|---|---|---|---|
| CNAG_01340 | Yta7 | 15 | 39 | 31 | 228 | 12 | 21 | ATPase, bromodomain |
| CNAG_03962 | Mcm6 | 13 | 17 | 51 | 60 | 15 | 16 | DNA helicase |

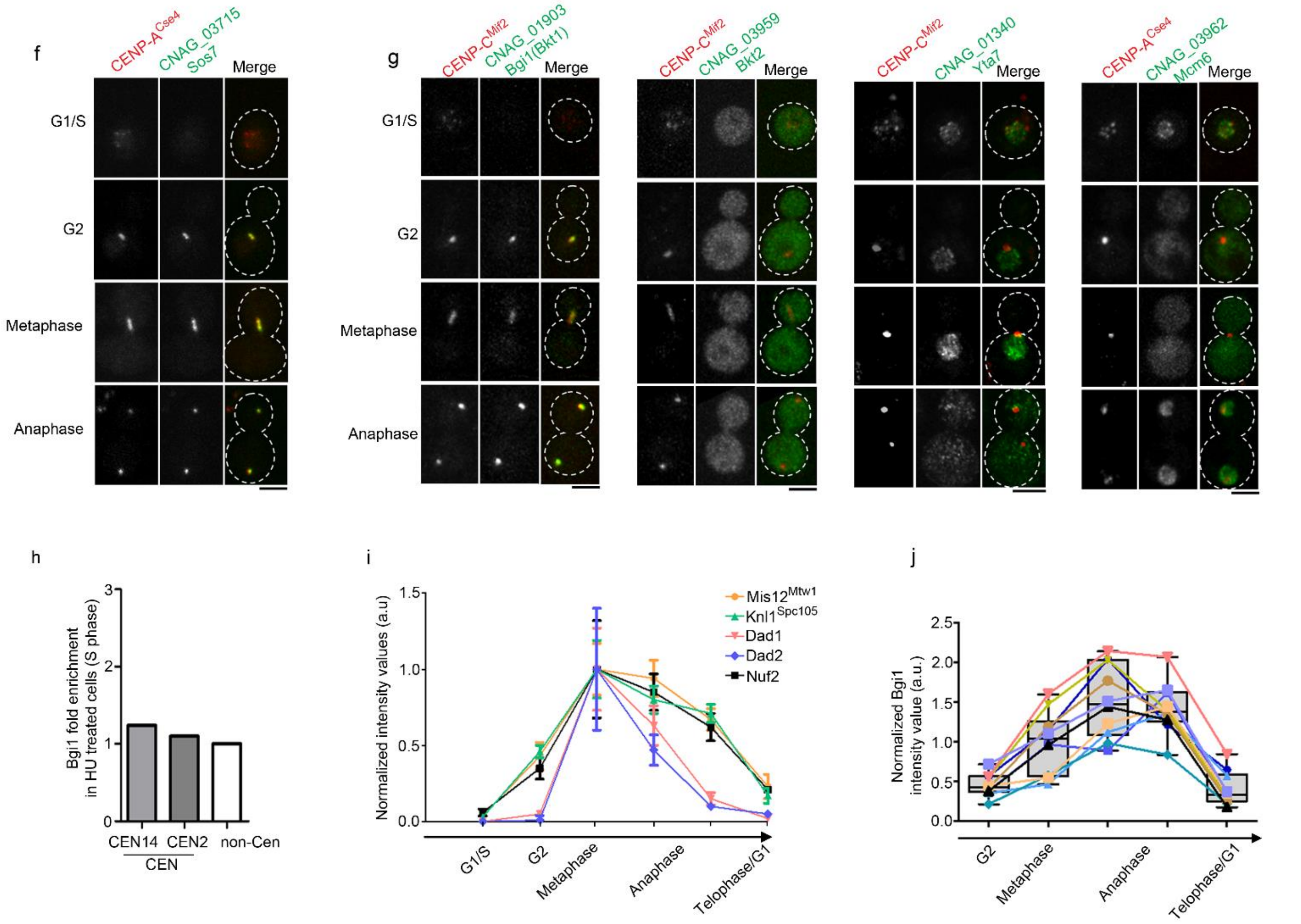


**Supplementary figure 1. Screening of obtained kinetochore interacting proteins. (a)** Immunoblot analysis of native 3xFLAG tagged kinetochore proteins. **(b)** Functional validation of 3xFLAG tagged kinetochore proteins by chromatin immuno-precipitation (ChIP-PCR). **(c)** Protein lysates were prepared from the mentioned strains. Kinetochore particles were purified using FLAG or GFP and analyzed by immunoblotting. **(d)** List of common interacting proteins with no known function. These protein hits showed conservation primarily across basidiomycetes and were identified in CENP-C$^{Mif2}$,

Dsn1, and Spc25 FLAG affinity-purified eluates. Predicted domains and possible domain functions are listed. **(e)** Tabulation of proteins known to bind chromatin with no strong association with kinetochore proteins and identified as interacting partners in CENP-C$^{Mif2}$, Dsn1, and Spc25 FLAG affinity purifications. **(f)** Sos7 (KNL1C) localization across cell-cycle stages. mCherry-CENP-A$^{Cse4}$ marks the kinetochore. Scale bar, 3 µm. **(g)** Localization of carboxy-terminus tagged GFP constructs across mentioned interphase and mitotic stages. CENP-C$^{Mif2}$-mCherry or CENP-A$^{Cse4}$-mCherry marks the kinetochores. Scale bar, 3 µm. **(h)** ChIP enrichment of bridgin in HU treated (200mM, 3hrs) cells arrested in early S phase. **(i)** Normalized signal intensities of Mis12$^{Mtw1}$-mCherry, Knl1$^{Spc105}$-GFP, GFP-Dad1, Dad2-mCherry and Nuf2-GFP across the interphase and mitotic stages. *N*=5. Error bars, s.d. **(j)** A normalized intensity plot of bridgin signals from G2 until telophase in ten cells. Error bars, standard deviation (s.d.).

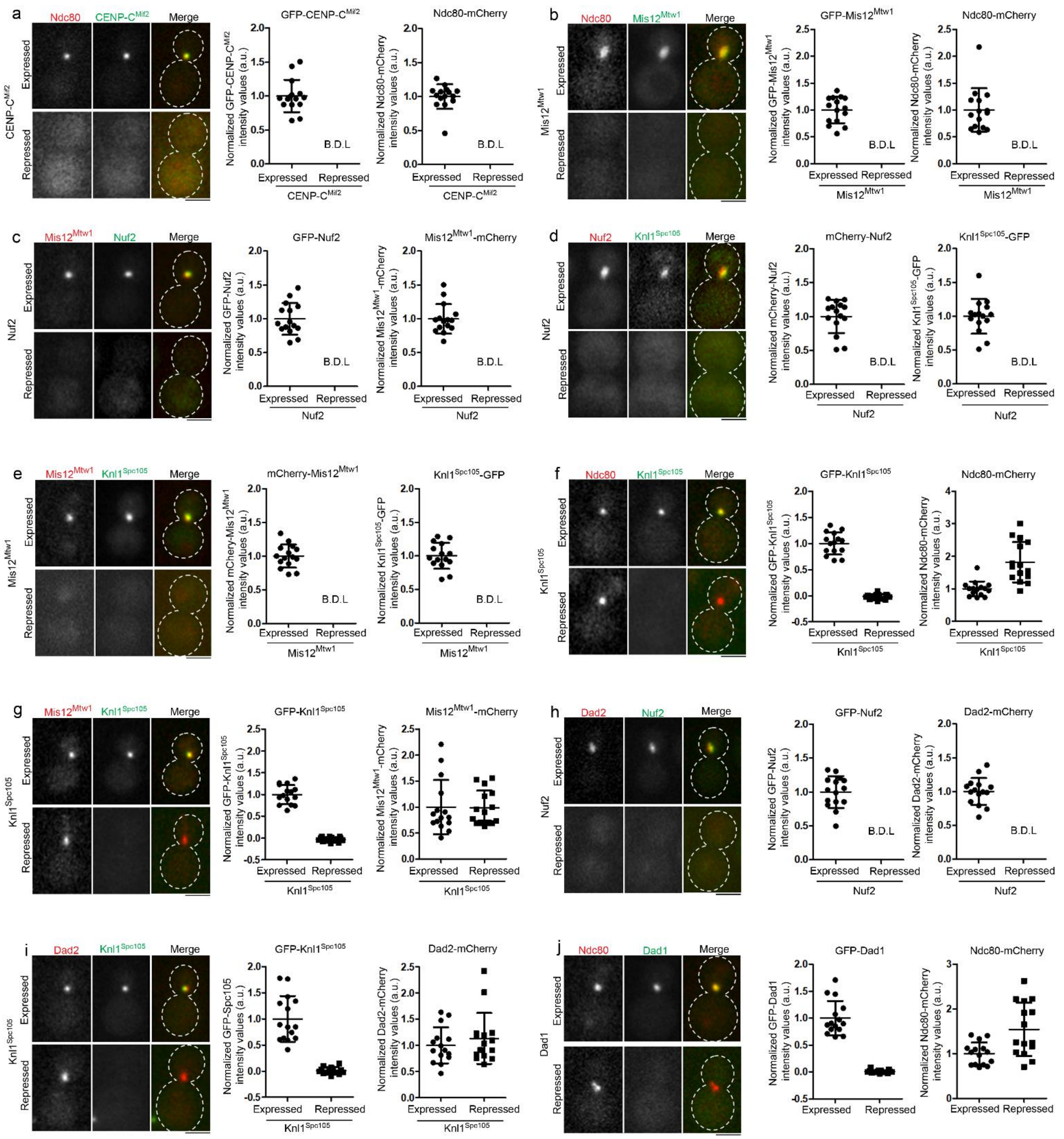


k

| | | | Depletion of kinetochore protein | | | | | | | |
|---|---|---|---|---|---|---|---|---|---|---|
| | Kinetochore sub-complex | | CENP-A | CENP-C | Mis12C | Ndc80C | KNL1C | Dam1 complex | | |
| | | KT protein | CENP-A$^{Cse4}$ | CENP-C$^{Mif2}$ | Mis12$^{Mtw1}$ | Nuf2 | Knl1$^{Spc105}$ | Dad1 | Dad2 | Ask1 |
| Kinetochore protein localization upon depletion | CENP-A | CENP-A$^{Cse4}$ | - | N.D. | + | + | + | + | N.D. | N.D. |
| | CENP-C | CENP-C$^{Mif2}$ | - | - | + | + | + | + | N.D. | N.D. |
| | Mis12C | Mis12$^{Mtw1}$ | - | - | - | - | + | + | N.D. | N.D. |
| | Ndc80C | Ndc80 | - | - | - | - | + | + | N.D. | N.D. |
| | KNL1C | Knl1$^{Spc105}$ | N.D. | N.D. | - | - | - | N.D. | + | N.D. |
| | Dam1 complex | Dad1 | - | N.D. | - | N.D. | N.D. | - | - | N.D. |
| | | Dad2 | - | - | N.D. | - | + | - | - | - |
| | | Ask1 | N.D. | N.D. | N.D. | N.D. | N.D. | N.D. | N.D. | - |

**(+)** : retains localization **(-)** : loss in localization N.D. : not determined

l

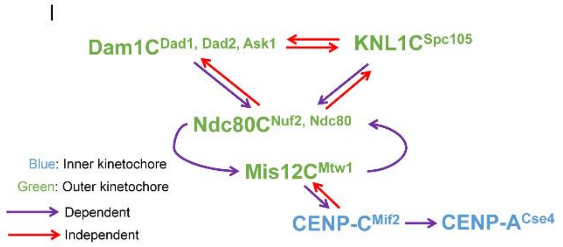

**Supplementary figure 2. Kinetochore localization interdependencies of protein sub-complexes at the *C. neoformans* kinetochore. (a-j)** Kinetochore proteins were fluorescently labeled in a kinetochore conditional mutant. As described in Fig 2b, normalized intensities of the tagged kinetochore proteins were measured under conditions of expression and repression of the conditional kinetochore mutant. The strong influence of the conditional kinetochore protein on the test protein resulted in signals that were below detectable levels (B.D.L). *N*=15 for the expressed and repressed conditions. Error bars, s.d. Scale bar, 3 µm. **(a)** The dependency of Ndc80 on CENP-C$^{Mif2}$. **(b)** The dependency of Ndc80 on Mis12$^{Mtw1}$. **(c)** The dependency of Mis12$^{Mtw1}$ (Mis12C) on Nuf2 (Ndc80C). **(d)** The dependency of Knl1$^{Spc105}$ (KNL1C) on Nuf2 (Ndc80C). **(e)** The dependency of Knl1$^{Spc105}$ (KNL1C) on Mis12$^{Mtw1}$ (Mis12C). **(f)** The dependency of Ndc80 (Ndc80C) on Knl1$^{Spc105}$ (KNL1C) **(g)** The dependency of Mis12$^{Mtw1}$ (Mis12C) on Knl1$^{Spc105}$ (KNL1C) **(h)** The dependency of Dad2 (Dam1C) on Nuf2 (Ndc80). **(i)** The dependency of Dad2 on Knl1$^{Spc105}$ (KNL1C) **(j)** The dependency of Ndc80 (Ndc80C) on Dad1 (Dam1C). **(k)** Table summarizing the kinetochore interdependencies tested. Symbols in red mark the shown interaction. **(l)** Schematic of interdependencies observed across kinetochore sub-complexes at the *C. neoformans* kinetochore.

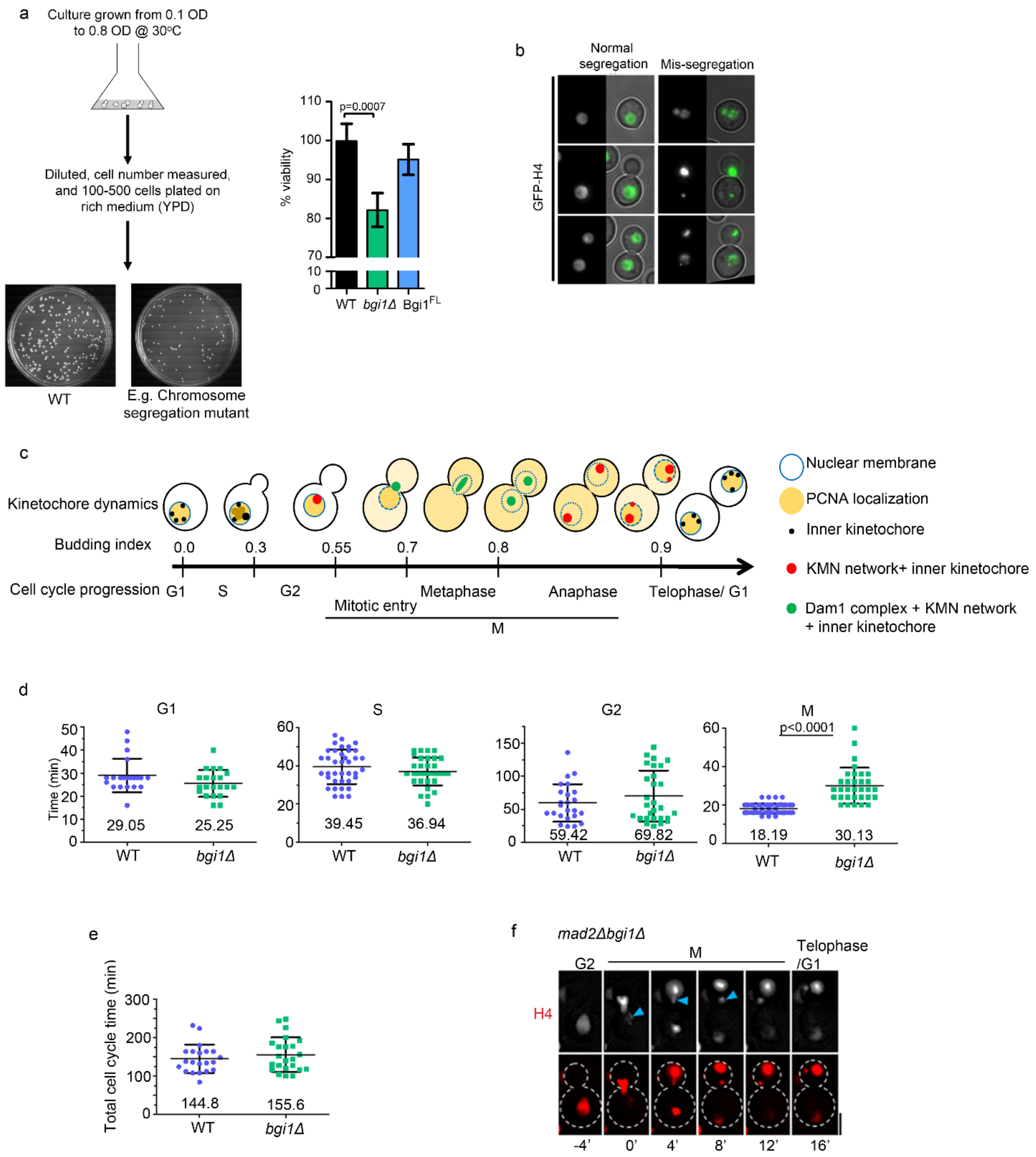


**Supplementary figure 3. Loss of bridgin triggers mis-segregation of chromosomes that leads to a reduced cell viability in *C. neoformans* cells. (a)** Schematic of experimental design to estimate cell viability *(left)*. CFU was counted after 48 h at 30ºC and tabulated *(right)*. Error bars, s.d. *N*=3. *P*-value determined using two-tailed t-test. **(b)** Representative events scored as normal or chromosome mis-segregation events are mentioned. Presence of multiple nuclei in a single cell, unequal segregation amongst daughter cells and formation of micronuclei were considered as mis-segregation events. **(c)** Graphical summary of cell cycle markers used to determine cell cycle stages in *C. neoformans* (*1*, *2*). H4 tagged strains was used where other markers were unavailable to determine stages of M phase. Budding index mentioned is an approximate daughter bud: mother bud diameter ratio observed in log phase growing cells. **(d)** Comparison of cell cycle stage-specific timing between WT and *bgi1Δ*. Mean of the measured times are mentioned. WT *N*=21,38,24 and 52 in G1, S, G2 and M phases respectively. *bgi1Δ N*=21, 34, 28 and 32 in G1, S, G2 and M phases respectively. Error bars, s.d. *P* value determined using two-tailed t-test. **(e)** Total cell cycle time was determined by live-cell analysis using cell cycle events marked by a nuclear protein PCNA and a chromatin-associated protein H4. Error bars, s.d. WT: *N*=21, *bgi1Δ*: *N*=23. *P* value determined using two-tailed t-test. **(f)** Representative image of *mad2Δbgi1Δ* time-lapse depicting formation of a daughter cell with micronuclei. Blue arrows point to an initial unattached chromosome that results in a micronuclei formation. Scale, 3 μm.

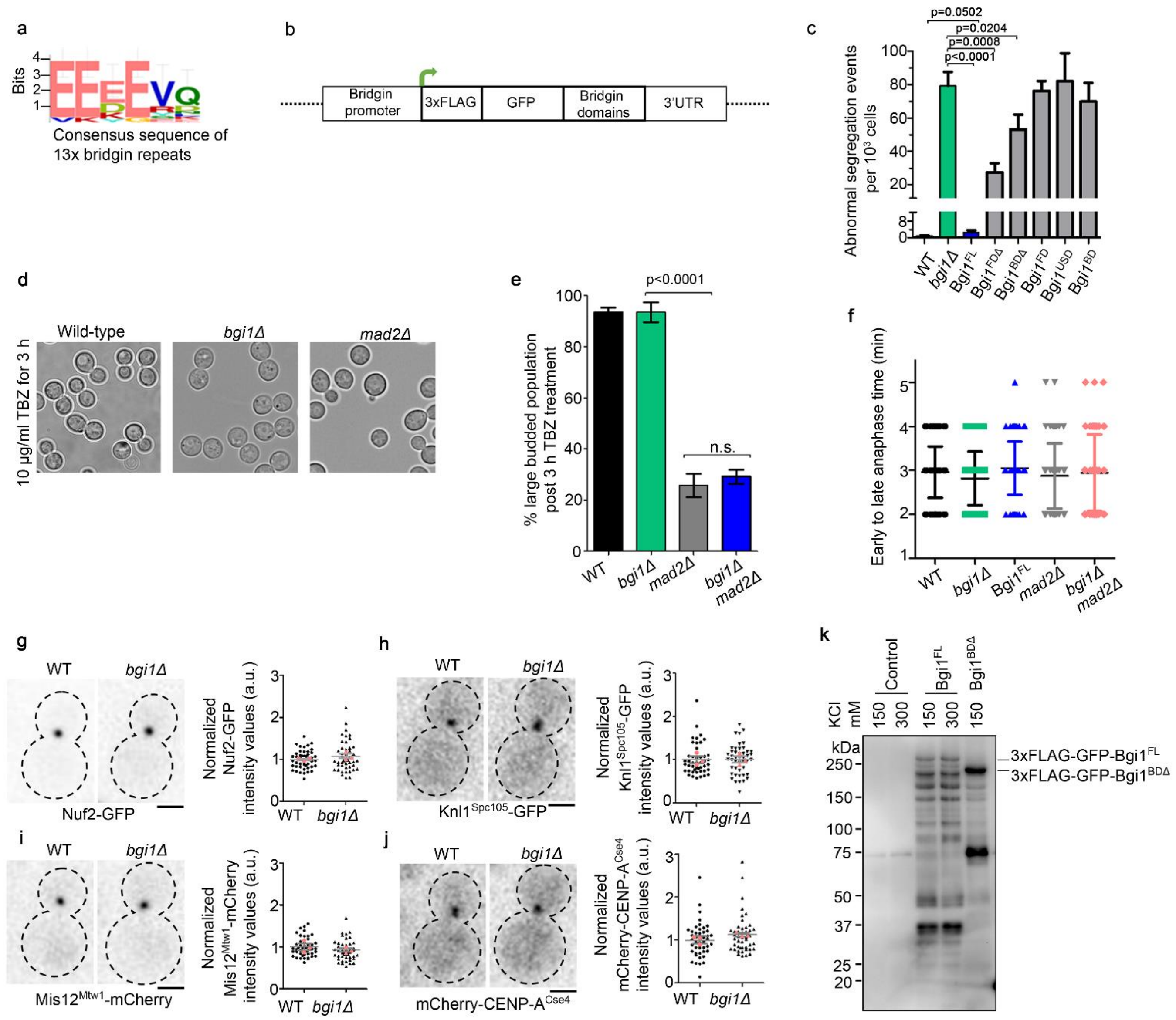


**Supplementary figure 4 Bridgin does not influence spindle assembly checkpoint function, spindle dynamics, and gross kinetochore composition. (a)** The motif for bridgin repeats was identified using MEME suit with the alignment of the 13 identified bridgin repeats, of 8-amino acid in length. **(b)** Schematic of the bridgin domain deletion constructs. **(c)** Rate of abnormal segregation defect was measured for each of the strains mentioned and normalized to events per $10^3$. A number of cells examined were >2000. *N*=3. Error bars, s.d. *P*-value determined using two-tailed t-test. **(d and e)** WT, *bgi1Δ,* and *mad2Δ* cells were treated for 3hrs with 10 µg/ml of thiabendazole (TBZ). **(d)** Representative bright-field micrographs of WT, *bgi1Δ,* and *mad2Δ* cells. **(e)** Percentage of large budded cells was determined by scoring for cells with a budding index of >0.55. Error bars, s.d. *N*>300 cells for each of the strains used. *P*-value determined using two-tailed t-test. **(f)** Measurement of time spent in anaphase for WT, *bgi1Δ*, *mad2Δ*, and *mad2Δbgi1Δ* cells. Cells were considered to have entered anaphase if their kinetochore distances were >1 µm or if the leading edge of the chromatin marker H4 were >1 µm. Late anaphase was defined as stages when kinetochore distances reached a maximum. *N*=88, 64, 61, 67 and 56 respectively for WT, *bgi1Δ*, $Bgi1^{FL}$, *mad2Δ* and *mad2Δbgi1Δ*. Error bars, s.d. **(g-j)** Measurement of the kinetochore protein intensity in WT and *bgi1Δ* cells. The kinetochore intensity values in representative cells of WT and *bgi1Δ* are shown as inverted grey scale images. Signal intensity was measured in 45 cells across 3 independent transformants. Red symbols denote the means of each independent transformant. Scale bar, 2 µm. *P* value determined using two-tailed t-test. **(g)** Nuf2-GFP**(h)** $Knl1^{Spc105}$-GFP **(i)** $Mis12^{Mtw1}$-mCherry **(j)** mCherry-CENP-$A^{Cse4}$. **(k)** Immunoblot analysis of 3xFLAG tagged bridgin constructs.

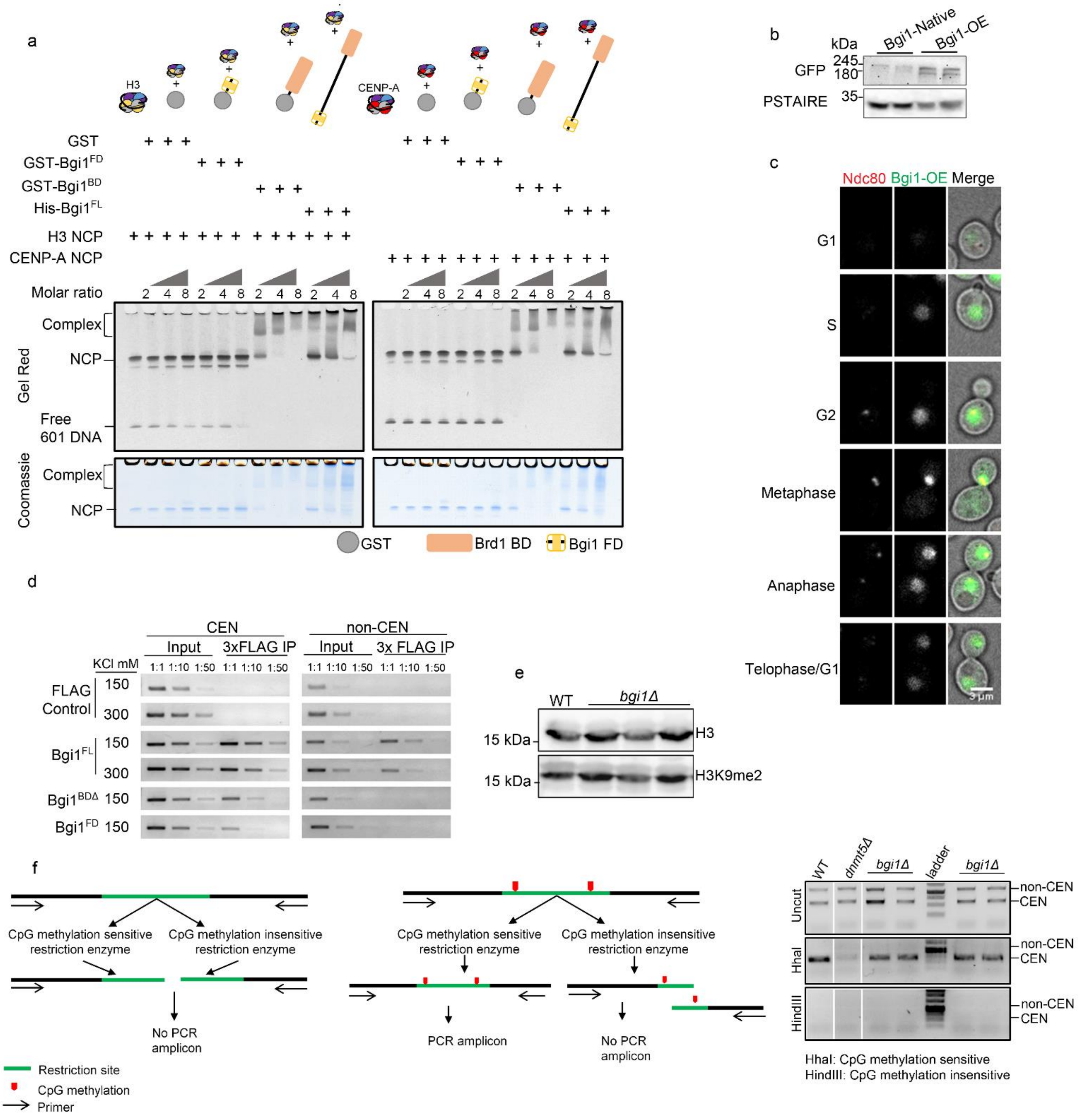


**Supplementary figure 5. The basic domain of bridgin can interact with nucleosomes, but it does not influence centromeric histone or centromere DNA methylation. (a)** EMSA performed with reconstituted GgH3 and HsCENP-A nucleosome core particles (NCP). Approximately 1 µM of reconstituted nucleosomes was incubated with the mentioned molar ratio of purified protein for 1 h at 4˚C. Samples were separated on a PAGE gel and stained with Gel Red followed by Coomassie. *N*=1. **(b)** The total cellular pool of bridgin was analyzed by immunoblot analysis. **(c)** Localization of GFP-Bgi1-OE at various cell cycle stages. Kinetochores are marked by the outer kinetochore protein Ndc80. **(d)** Native ChIP of 3xFLAG-GFP bridgin protein derivatives. G2/M cells were enriched and lysed using bead beating. FLAG affinity purification was performed as in Fig 5a**.** DNA was isolated from the elute and PCR was set with two centromere and non-centromere primers. Dilutions of eluted DNA and input of 1:1, 1:10 and 1:50 were used. *N*=1. **(e)** Immunoblot analysis of the whole-cell pool of H3K9me2 levels in WT and *bgi1Δ* cells. Cell lysates from an asynchronous culture of WT and *bgi1Δ* were prepared by bead beating. **(f)** Detection of CpG methylation by Dnmt5 in WT and *bgi1Δ*.

Schematic of the restriction enzyme-PCR based assay used to assess the methylation status at the centromere *(left)*. Restriction enzyme HindIII is insensitive, while HhaI is sensitive to CpG methylation. Genomic DNA was isolated and digested with either HindIII or HhaI and primers flanking the restriction site is used to estimate relative levels of digested genomic DNA, a read-out for levels of CpG methylation at the locus. Amount of HhaI PCR amplicon is proportional to the level of CpG methylation, in comparison to WT.

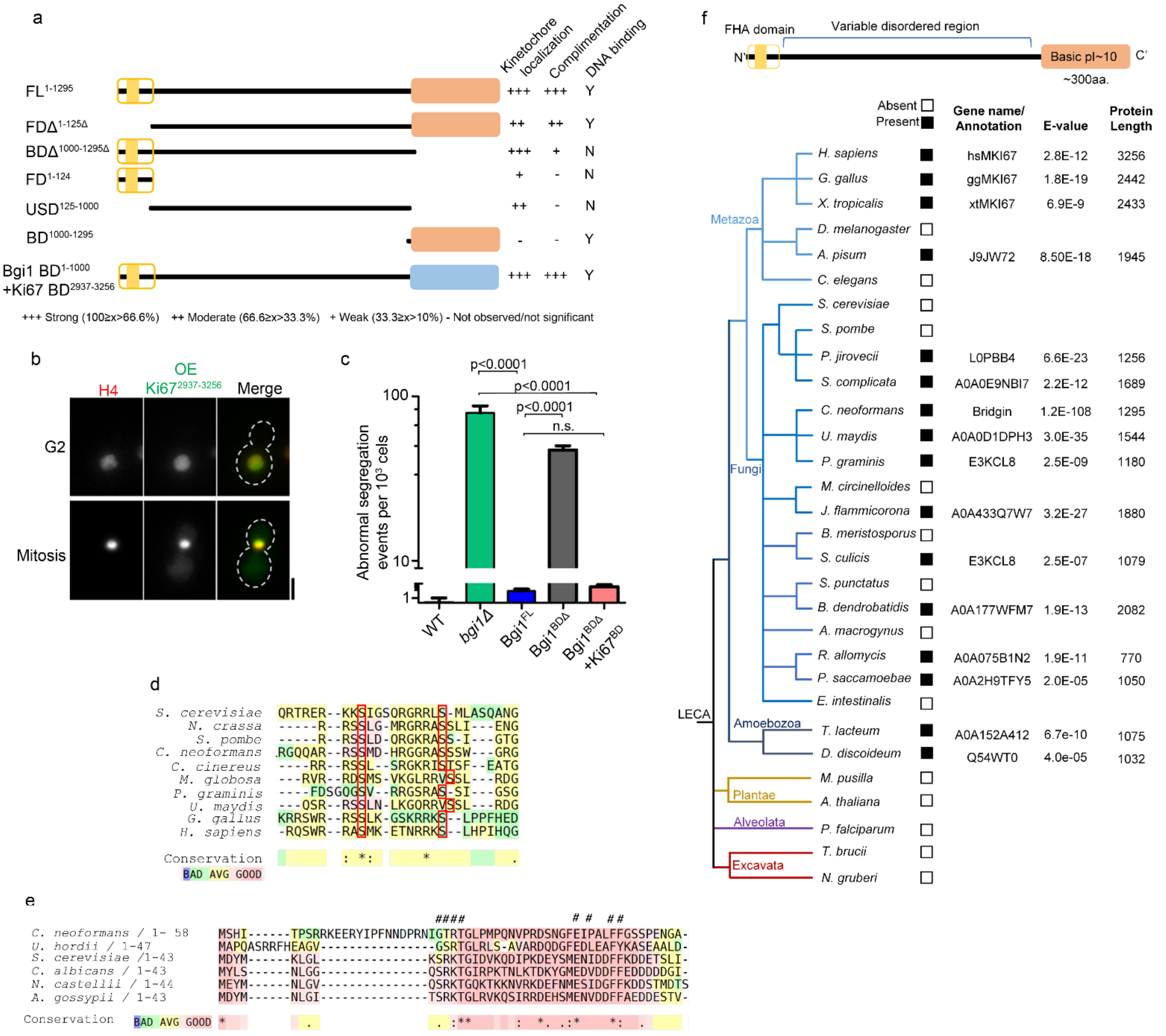


**Supplementary figure 6. Homologs of the novel linker protein bridgin were identified outside the fungal kingdom. (a)** Summary of functional analysis of the bridgin protein derivatives truncated for various domains based on their kinetochore localization, functional complementation, and ability to bind DNA *in vivo.* **(b)**Localization of GFP-Ki67$^{BD}$-OE at G2 and mitosis. Chromatin is marked using H4-mCherry. Scale bar, 3 µm. **(c)** The extent of complementation by Bgi1$^{FL}$, Bgi1$^{BD\Delta}$ or basic domain swap bridgin chimeric constructs was measured. The number of mis-segregation events per 1000 cells was estimated. Cells were grown to early log phase (0.8-1 OD) at 30˚C and abnormal segregation events were scored using the chromatin marker H4-mCherry. Error bars, s.d. >1000 for each mentioned strain were measured. *N*=3. *P*-value determined using two-tailed t-test. **(d)** Sequence alignment of the Dsn1 basic motif encompassing the two Aurora kinase B/Ipl1 phosphorylation sites across species are highlighted in the red box. Alignment and visualization were performed using T-coffee. **(e)** Alignment of the described CENP-C$^{Mif2}$-Mis12$^{Mtw1}$ interacting motif in CENP-C$^{Mif2}$. # represent CENP-C$^{Mif2}$ residues important for CENP-C$^{Mif2}$-Mis12$^{Mtw1}$ interaction, as shown in *S. cerevisiae*. **(f)** Identification of bridgin-like proteins across eukaryotes. Parameters scored for include the FHA domain within the first ~200 amino acids, followed by a variable-length disordered region and a carboxy terminus of ~300 residues with an isoelectric point of ~10 or greater are considered.
